# Typical resting state activity of the brain requires visual input during an early sensitive period

**DOI:** 10.1101/2021.06.09.446724

**Authors:** Katarzyna Rączy, Cordula Hölig, Maria J. S. Guerreiro, Sunitha Lingareddy, Ramesh Kekunnaya, Brigitte Röder

## Abstract

Sensory deprivation, following a total loss of one sensory modality e.g. vision, has been demonstrated to result in intra- and cross-modal plasticity. It is yet not known to which extent intra- and cross-modal plasticity as a consequence of blindness reverse if sight is restored.

Here, we used functional magnetic resonance imaging to acquire blood oxygen level dependent (BOLD) resting state activity during an eyes open and an eyes closed state in congenital cataract-reversal individuals, developmental cataract-reversal individuals, congenitally permanently blind individuals and sighted controls. The amplitude of low frequency fluctuations (ALFF) of the BOLD signal - a neural marker of spontaneous brain activity during rest - was analyzed.

As has been shown before, in normally sighted controls we observed an increase in ALFF during rest with the eyes open compared to rest with eyes closed in visual association areas and in parietal cortex but a decrease in auditory and sensorimotor regions. In congenital cataract-reversal individuals, we found an increase of the amplitude of slow BOLD fluctuations in visual cortex during rest with eyes open compared to rest with eyes closed too but this increase was larger in amplitude than in normally sighted controls. At the same time, congenital cataract-reversal individuals lagged a similar increase in parietal regions and did not show the typical decrease of ALFF in auditory and sensorimotor cortex. Congenitally blind individuals displayed an overall higher amplitude in slow BOLD fluctuations in visual cortex compared to sighted individuals and compared to congenital cataract-reversal individuals in the eyes closed condition.

Higher ALFF in visual cortex of congenital cataract-reversal individuals than in normally sighted controls during eyes open might indicate an altered excitatory-inhibitory balance of visual neural circuits. By contrast, the lower parietal increase and the missing downregulation in auditory and sensorimotor regions suggest a reduced influence of the visual system on multisensory and the remaining sensory systems after restoring sight in congenitally blind individuals. These results demonstrate a crucial dependence of multisensory neural networks on visual experience during a sensitive phase in human brain development.

## Introduction

A number of studies have demonstrated that sensory deprivation, due to e.g. blindness, results in functional and structural changes of the brain related to intra- and cross-modal plasticity.^1,2^ While intramodal plasticity comprises reorganizations of neural systems predominantly linked to the intact sensory modalities, e.g. the auditory system,^3-5^ cross-modal plasticity refers to an activation of neural circuits primarily associated with the deprived sensory modality, e.g. the visual cortex, by input of the intact sensory modalities (e.g. auditory input). A higher activation of visual cortex in permanently blind humans has been found during rest,^6,7^ as well as during perceptual (e.g. tactile discrimination, auditory localization, motion perception) and higher cognitive tasks (e.g. Braille reading, verbal memory tasks, spoken sentence and arithmetic processing).^8,9^ Both intra- and cross-modal plasticity have often been associated with compensatory performance in blind humans.^10^

The question has been raised whether adaptations to blindness, in particular cross-modal plasticity interfere with functional recovery when vision is restored.^2,11,12^ So far no clear evidence has been provided to support this notion.^2,13,14^ While a larger influence of the auditory system on the visual system has been demonstrated in sight recovery individuals with short deprivation epochs using brain imaging^15-18^ and behavioral measures,^19^ electrophysiological studies have not yet found analogous evidence.^14^ Recent results in the deaf cat have argued against cross-modal plasticity limiting auditory recovery.^20,21^ Crucially, cross-modal activation in sight recovery individuals’ visual cortex^15,16^ was rather weak compared to the typical cross-modal activation observed in permanently blind humans.^22^ However, these results were from only shortly deprived sight recovery individuals and thus, longer deprivation periods might cause more extensive and persisting cross-modal plasticity. Alternatively, it has been suggested that what causes persisting visual deficits in multiple visual functions^23,24^ is the lack of a functional and structural tuning of genuine visual processing as which was suggested by non-human animal studies to happen under the control of visual experience during early brain development.^25,26^

Resting brain activity is considered as crucial scaffold for task related processing. For example, it has been found that resting state connectivity in the visual cortex of awake ferrets reflects a functional organization of the brain according to the expected visual input.^27^ In humans, resting state blood oxygen level dependent (BOLD) fluctuations - predominantly in the low-frequency range (0.01–0.1 Hz, i.e. low frequency oscillations) - have been likewise shown to correlate across functionally related, even spatially remote brain regions.^28^ These correlations of spontaneous BOLD fluctuations between brain regions are known as resting state functional connectivity and have been proposed to reflect distinct brain networks.^29-31^ For example, Biswal et al.^28^ demonstrated that low frequency BOLD fluctuations (0.01-0.08 Hz) within the sensorimotor network during rest mirrored typical sensorimotor activity during a finger tapping task. Resting state BOLD connectivity often reflects structural connectivity as assessed with DTI in humans (diffusion tensor imaging)^32^ and as demonstrated by combining noninvasive fMRI and invasive anatomical retrograde tracing methods in monkeys.^33^ Resting state connectivity, does not, however, provide information about the level of BOLD signal change. In contrast, the Amplitude of Low Frequency Fluctuations (ALFF)^34^ - which is defined as the total power within the frequency range between 0.01 and 0.08 Hz - has been demonstrated to indicate the intensity of BOLD fluctuations and thus the overall neural activity level.

In the normally sighted population ALFF has been contrasted between eyes open and eyes closed resting states. These studies have observed significantly increased ALFF during eyes open relative to eyes closed in the bilateral middle occipital gyrus (BA19/39) and in orbital frontal cortex (BA47/11), whereas a decreased ALFF was found in the bilateral pre- and postcentral gyrus (BA2/3, 4, 6), as well as in temporal (BA21, 22) and insula regions (BA22/48), the thalamus, and in the calcarine sulcus (BA17).^35-37^ ALFF has additionally been used to document characteristic deviations in resting state activity in psychiatric and neurological populations.^38-44^

In simultaneous EEG-fMRI studies a negative correlation of alpha oscillatory activity and resting state BOLD connectivity within the visual cortex has been reported in an eyes opened vs. eyes closed comparison^45^ and during resting state with eyes closed.^46,47^ Such negative correlations were observed in particular for BOLD connectivity between the primary visual cortex and other occipital brain regions^48^ and were specific for the alpha band. Alpha band activity has been suggested to reflect a neural mechanism important for the control of the excitatory-inhibitory balance of neural circuits to guarantee efficient processing: During task processing alpha band power is high for neural circuits not engaged in task processing but low for task relevant neural systems^49^ resulting in an improved signal-to-noise ratio in neural networks.

Posterior alpha oscillatory activity was found to be significantly reduced in congenital cataract-reversal individuals as compared to developmental cataract-reversal individuals (that is, individuals with a history of late onset cataracts) and typically sighted controls.^50^ The authors speculated that the reduced alpha activity in congenital cataract-reversal individuals was due to a reduction of inhibitory mechanisms which seem to be elaborated during sensitive periods.^26^ In fact, reduced posterior alpha activity is a typical characteristic of the EEG of permanently blind humans.^51-54^ Moreover, two MEG studies later reported enhanced gamma activity in congenitally permanently blind individuals compared to typically sighted controls.^55,56^ Recent studies have suggested that alpha and gamma oscillatory activity indicate antagonistic mechanisms with alpha oscillations controlling gamma oscillatory activity.^57,58^ Interestingly, in monkeys it has been observed that slow BOLD fluctuations correlated with high frequency local field potentials (LFP) in the gamma range particularly in an eyes closed condition.^59^

Based on the group differences in oscillatory activity (lower alpha activity in both congenital cataract-reversal and congenitally blind individuals, higher gamma activity in congenitally blind humans) we hypothesize that ALFF is enhanced in congenital cataract-reversal individuals as well as in congenitally blind individuals compared to typically sighted controls in occipital brain regions. However, resting state brain activity has not yet been investigated in congenital cataract-reversal individuals. Since resting state activity builds the foundation of stimulus driven activity and might reflect a reinforcing process for the existing functional connectivity,^59,60^ assessing the amplitude of low frequency oscillations of the BOLD signal in congenital cataract-reversal individuals compared on the one hand, to congenitally permanently blind humans, and on the other hand, to typically sighted controls allows investigating the degree of functional recovery of visual neural circuit functioning following sight restoration. Hence, the present study allowed investigating the question of whether there is a sensitive period for the development of typical resting state activity in humans.

Resting state fMRI (rsfMRI) was recorded both during eyes open and eyes closed in four groups: (1) congenital cataract-reversal individuals (CC group) who had experienced visual deprivation since birth for up to 18 years due to bilateral dense congenital cataracts, (2) congenitally permanently blind individuals (CB group) to indicate the adaptation of resting state activity to congenital blindness, (3) developmental cataract-reversal individuals (DC group) who had developed cataracts later during childhood to serve as a control for visual impairments and other effects related to a history of cataracts and cataract removal surgery and (4) typical sighted controls (SC group).

Low frequency oscillations of the BOLD signal were assessed as described in Yu-Feng et al.^34^ Based on previous EEG results (reduction of alpha oscillatory activity in CC and CB individuals, higher gamma band activity in CB individuals as discussed above) we hypothesized to find higher ALFF in visual areas of CC and CB individuals compared to the SC group. As previous studies have revealed retracted cross-modal activity in the CC group^16,61^ compared to what has been typically found in CB individuals,^2,9^ we predicted on the one hand that ALFF is lower in the CC group than in the CB group in early visual areas and on the other hand that ALFF is modulated by EO vs. EC in the CC group similarly as seen in normally sighted individuals. In particular, we hypothesized to replicate the typical decrease of ALFF in the eyes open compared to the eyes closed condition in early visual cortex and an increase in higher occipital regions in both the CC and the SC group. Since previous research on resting state connectivity in CB individuals indicated lower connectivity of visual and both sensorimotor and auditory areas,^62^ we predicted a lower decrease of ALFF for the EO compared to the EC condition in these areas in the CC group compared to the SC group. We expected the CC vs. SC group difference to be specific for the CC group, that is, we predicted to not finding the same pattern of results for the DC vs. SC group comparison.

## Materials and Methods

### Participants

All participants were recruited at the LV Prasad Eye Institute (LVPEI) and from the local community of Hyderabad (India). Four groups were included: congenital cataract-reversal individuals (CC group), developmental cataract-reversal individuals (DC group), congenitally permanently blind individuals (CB group) and sighted controls (SC group).

The original CC group consisted of 22 individuals. Three of them were excluded from the final analysis; one because of insufficient data due to premature termination of the scanning session (C7), a second due to a deprivation period of less than three months (C12) and in a third participant (C13) it turned out later that cataracts had not been dense. The final CC group consisted of 19 individuals (nine males, 10 females, mean age: 16.9 years, range: 6-36 years, mean age at surgery: 67.9 months, range: 3–216 months; mean time since surgery: 138.9 months, range: 6-412 months). Mean logMAR visual acuity in the better eye (based on the most recent entry in the medical records) was 0.86 (range: 0.30–1.80). One participant’s visual acuity could not be measured with the letter charts. Thus visual acuity was assessed with the maximal distance at which the participant was able to count fingers, which in this particular case was 1 m (which was translated to 1.80 logMAR). The history of bilateral dense congenital cataracts was confirmed based on the information available in the medical records. In the classification process, additionally to the clinical diagnosis, we considered factors such as presence of sensory nystagmus and strabismus, absence of fundus view prior to surgery, and a positive family history. For detailed information about the participants, see Table 1.

**Table 1.** Clinical and Demographic Information for Congenital Cataract-Reversal Participants.

| Participant | Sex | Age (years) | Cataract onset | Pre-surgery visual acuity in the better eye (logMAR) | Age at surgery (months) | Visual acuity in the better eye at testing (logMAR) | Additional details |
| --- | --- | --- | --- | --- | --- | --- | --- |
| CC1 | M | 24 | Congenital | N/A | 5 | 0.80 | Nystagmus, esotropia |
| CC2 | M | 36 | Congenital | N/A | 24 | 0.40 | Nystagmus, esotropia |
| CC3 | M | 9 | Congenital | CF at 3m<sup>a</sup> | 84 | 0.80 | Nystagmus, |
| CC4 | F | 6 | Congenital | PL+, PR+ | 48 | 0.90 | Nystagmus, |
| CC5 | F | 18 | Congenital | CFCF<sup>a</sup> | 192 | CF at 1m (1.80 logMAR) | Nystagmus, exotropia, microcornea - |
| CC6 | M | 32 | Congenital | N/A | 72 | 1.30 | Nystagmus, |
| CC7* | M | 11 | Congenital | PL+, PR+ | 61 | 0.78 | Nystagmus, esotropia |
| CC8 | F | 28 | Congenital | CFCF | 216 | 1.10 | Nystagmus, esotropia |
| CC9 | M | 11 | Congenital | FFL | 5 | 0.90 | Nystagmus, esotropia |
| CC10 | M | 28 | Congenital | N/A | 168 | 1.00 | Nystagmus, |
| CC11 | M | 13 | Congenital | FFL | 15 | 0.30 | Nystagmus, exotropia |
| CC12* | M | 8 | Congenital | FFL | 1 | 0.20 | Pseudophakia<sup>b</sup> |
| CC13* | M | 30 | Congenital | 1.10<sup>a</sup> | 216 | 1.30 | Pseudophakia, Nystagmus |
| CC14 | F | 26 | Congenital | N/A | 8 | 0.80 | Exotropia<sup>b</sup> |
| CC15 | F | 10 | Congenital | FFL | 42 | 0.40 | Nystagmus, exotropia |
| CC16 | F | 7 | Congenital | FFL | 3 | 0.80 | Nystagmus, esotropia |
| CC17 | F | 17 | Congenital | CFCF<sup>a</sup> | 123 | 1.48 | Nystagmus<sup>c</sup> |
| CC18 | F | 11 | Congenital | CF at 1m | 127 | 1.30 | Nystagmus |
| CC19 | M | 8 | Congenital | FFL<sup>a</sup> | 64 | 0.40 | Nystagmus, esotropia |
| CC20 | F | 18 | Congenital | FFL at 1 m | 25 | 0.48 | Nystagmus, esotropia, microcornea |
| CC21 | M | 13 | Congenital | CF at 0.5m | 64 | 0.74 | Nystagmus, esotropia |
| CC22 | F | 6 | Congenital | No FFL | 5 | 0.60 | Nystagmus, esotropia |

Note. M = male; F = female; N/A = not available; OU = both eyes; OS = left eye; CF = counting fingers at *n* meters; PL+ = able to perceive light; CFCF = counting fingers close to face; HM+ = able to perceive hand motion; FFL = fixing and following light; PR+ = able to report the location of light.
<sup>a</sup> partially absorbed cataracts
<sup>b</sup> the presence of nystagmus was not reported in the medical file
<sup>c</sup> operated only in one eye
\* excluded participants

The original DC group consisted of 16 individuals. Five participants (D12-16) were excluded from the final analysis. We were not able to come to an unambiguous classification and thus refrained from including these participants in either the DC group or the CC group. The final DC group consisted of 11 individuals (eight males, three females, mean age: 15.8 years, range: 9-43 years, mean age at surgery: 157.3 months, range: 84–484 months; mean time since surgery: 37.0 months, range: 7-60 months; mean logMAR visual acuity, according to the most recent entry in the medical records: 0.37, range: 0.10–1.78; For detailed information about the participants, see Table 2). Note that though the age at surgery is known in DC individuals, the exact age at cataract onset is hard to define, since developmental cataracts typically gradually emerge. Furthermore, developmental cataracts were not necessarily dense as in the CC group.

**Table 2.** Clinical and Demographic Information for Developmental Cataract-Reversal Participants.

| Participant | Sex | Age (years) | Cataract onset | Pre-surgery visual acuity in the better eye (logMAR) | Age at surgery (months) | Visual acuity in the better eye at testing (logMAR) | Additional details |
| --- | --- | --- | --- | --- | --- | --- | --- |
| DC1 | M | 13 | Developmental | PL+, PR+ | 108 | 1.78 | Nystagmus, exotropia OU |
| DC2 | M | 13 | Developmental | 0.80 | 150 | 0.10 | — |
| DC3 | F | 43 | Age-related<sup>b</sup> | 0.60 | 484<sup>c</sup> | 0.30 | — |
| DC4 | M | 12 | Congenital | 0.80 | 142 | 0.60 | — |
| DC5 | M | 9 | Developmental | 1.04 | 86 | 0.30 | — |
| DC6 | M | 9 | Developmental | 0.90 | 84 | 0.10 | — |
| DC7 | M | 17 | Developmental | 0.40 | 150 | 0.10 | — |
| DC8 | M | 13 | Developmental | 1.00 | 96 | 0.20 | Exotropia |
| DC9 | M | 11 | Developmental | 0.48 | 100 | 0.30 | — |
| DC10 | F | 13 | Developmental | 0.50 | 122 | 0.10 | — |
| DC11 | F | 21 | Developmental | 0.40 | 208 | 0.20 | — |
| DC12* | F | 34 | Congenital | 1.48<sup>a</sup> | 376 | 1.10 | Nystagmus, exotropia |
| DC13* | M | 32 | Developmental | CF at 2 m | 108 | 1.30 | Nystagmus, iris coloboma OU |
| DC14* | M | 6 | Congenital | 1.30 | 67 | 0.80 | Nystagmus |
| DC15* | M | N/A | Congenital | N/A | N/A | 0.40 | — |
| DC16* | F | N/A | Congenital | CF at 2 m<sup>a</sup> | 183 | 1.30 | Exotropia |

Note. M = male; F = female; OU = both eyes; CF = counting fingers at *n* meters; PL+ = able to perceive light; PR+ = able to report the location of light.
<sup>a</sup> partially absorbed cataracts
<sup>b</sup> cataracts developed after the age of twelve
<sup>c</sup> operated only in one eye
\* excluded participants

The DC group served as a control for visual impairments and other effects related to a history of cataracts and cataract removal surgery.

The CB group consisted of 12 congenitally blind individuals. Three participants were excluded from the final analysis; one (CB6) due to the lack of the eyes open condition, a second (CB11) due to a central cause of blindness and a third (CB12) due to non-congenital blindness. The final CB group consisted of nine individuals (six males, three females, mean age: 20 years, range: 9-39 years). For detailed information about the participants, see Table 3.

**Table 3.** Clinical and Demographic Information for Congenitally Blind Participants.

| **Participant** | **Sex** | **Age**<br>**(years)** | **Cause of blindness / Diagnosis** | **Blindness**<br>**onset** | **Visual**<br>**acuity** |
| --- | --- | --- | --- | --- | --- |
| CB1 | M | 39 | Microphthalmia OU | Congenital | N/A |
| CB2 | M | 9 | Lebers congenital amaurosis OU | Congenital | FFL |
| CB3 | M | 21 | Microphthalmos OU, Microcornea OU, | Congenital | NLP |
| CB4 | F | 19 | Microphthalmos OU | Congenital | PL+ |
| CB5 | M | 17 | Phthisis Bulbi OD, Anterior Staphyloma OS | Congenital | PL+ |
| CB6* | M | 16 | Microphthalmos OD, Anophthalmos OS | Congenital | PL+ |
| CB7 | F | 19 | Anterior staphyloma OD, Phthisis Bulbi OS | Congenital | PL+ |
| CB8 | M | 23 | Corneal scar OU, Microphthalmia OU | Congenital | CFCF |
| CB9 | M | 16 | Microphthalmia OU, band-shaped Keratopathy OU | Congenital | PL+, PR+ |
| CB10 | F | 17 | Lebers Congenital Amaurosis OU | Congenital | NLP |
| CB11* | M | N/A | Cortical lesions | Congenital | PL+ |
| CB12* | N/A | N/A | N/A | Late-onset | N/A |

Note. M = male; OU = both eyes; OD = right eye; OS = left eye; N/A = not available; FFL = fixing and following light; NLP = no light perception; F = female; PL+ = able to perceive light; PR+ = able to report the location of light; CFCF = counting fingers close to face.
\* excluded participants

The SC group comprised 28 individuals (19 males and nine females, mean age: 21.6 years, range: 6-56 years). Nineteen out of 28 sighted controls were matched in age and sex with the CC group (11 males and eight females, mean age: 19.8 years, range: 6-41 years). Eleven out of 28 sighted controls were matched in age and sex with the DC group (eight males and three females, mean age: 16.8 years, range: 10-42 years). Nine out of 28 sighted controls were matched in age and sex with the CB group (six males and three females, mean age: 20.8 years, range: 10-41 years). All sighted control participants had normal or corrected to normal vision.

All participants or their legal guardians (in case of minors) provided written informed consent and an assent (in case of minors) prior to taking part in the study. The subjects’ consent was obtained according to the Declaration of Helsinki. Participants and in case of minors, their legal guardians received a small compensation for the time of participation (e.g. lost wages) and for other expenses such as travel costs.

The study was approved by the Local Ethics Board of the Faculty of Psychology and Movement Sciences, University of Hamburg, Germany, the Ethics Board of the German Psychological Society as well as the Institutional Ethical Review Board of the LVPEI.

### MRI acquisition

Data were acquired at a radiology clinic (Lucid Medical Diagnostics Banjara Hills, Hyderabad, India), with a 1.5 T GE Signa HDxt scanner. Resting state functional MR scans were collected employing a gradient-echo echo-planar imaging (EP/GR) sequence. An 8-channel head coil was used (flip angle = 90°; TR = 2000 ms; TE = 30 ms, FOV = 220 x 220 mm; 64 x 64 matrix). TRs varied slightly among the participants (range: 1950-2300) due to the participant’s head size, body weight and height. This TR adjustment was built into the available GE protocol. Thirty-two (or in one subject 38, in a second - 34 and in a third – 33) interleaved axial slices (thickness 3 mm; in-plane resolution = 3.4 x 3.4 mm^2^, interslice gap = 4 mm) in ascending order were acquired. Anatomical T1-weighted images, using 3D-spoiled gradient recalled (3D-SPGR) sequence (TR = 1470 ms (range: 1466-1503); TE = 6.62 ms (range: 6.60-6.76), FA = 15°; on average 187 axial slices (range: 172-196 slices); voxel dimensions = 0.8 × 0.4297 × 0.4297 mm; matrix size = 512 × 512; inversion time = 500ms, slice thickness = 1.6 mm, slice gap = -0.8 mm) were additionally acquired for each subject.

### MRI acquisition – Procedure

Two runs of resting state fMRI were acquired for each participant, one with eyes open (EO) and one with eyes closed (EC). Each run lasted for 8.53 minutes: 45 participants started with the EO condition and the remaining 22 with the EC condition. Note that counterbalancing of the conditions across groups was not perfect due to miscommunication in a clinical setting: 12 CC individuals started with the EO and seven with the EC, eight DC individuals started with the EO and three with the EC, seven CB individuals started with the EO and two with the EC, 18 SC individuals started with the EO and 10 with the EC. During the resting state fMRI scanning, in both conditions, the participants were asked to lay as still as possible, to not think about anything in particular and to not fall asleep. In the EO condition, the subjects were asked to keep their eyes open and in the EC condition to keep their eyes closed throughout the whole run. The scanner room was kept dimly lit during scanning.

### Data Preprocessing

Data preprocessing and analysis of ALFF was performed using the DPARSF software of DPABI V4.3.^63^ DPABI is a toolbox for Data Processing and Analysis of Brain Imaging based on SPM12 and REST implemented in MATLAB (MathWorks, Natick, MA, USA).

First, basic screening of images (visual inspection) for each participant was performed to check for image quality. Exclusion criteria were: signal loss due to susceptibility artifacts and cut off slices. No images were excluded due to this procedure. Then, all time points were transformed from EPI DICOM to NIFTI. The first 10 time points were removed for signal stability. Next, a standard pre-processing pipeline was applied: All the acquired functional volumes were aligned to the first slice for EPI distortion and slice acquisition time (slice timing). The functional volumes were subsequently spatially realigned (using rigid body transformations to correct for head movements), spatially normalized to the standard adult East Asian template (MNI space), and smoothed with a 4 mm (FWHM) Gaussian kernel. In the next step linear trends were removed from the time series (detrend) using polynomial regressors. Structural images were segmented into grey matter (GM), white matter (WM) and cerebrospinal fluid (CSF). Subsequently, nuisance covariates were regressed out: Friston 24 head motion parameters, CSF and WM signals. The use of full nuisance regression including polynomial detrending in ALFF data optimizes the group-level analysis.^64^ In the next step the time series for each voxel were filtered (band-pass filtering: 0.01–0.08 Hz),^28,36^ to remove the effects of very-low-frequency drifts and high frequency noise. For a given voxel, a fast Fourier transform (FFT) (parameters: taper percent=0, FFT length=shortest)^65^ was used to convert the filtered time series to the frequency domain to obtain the power spectrum. The power spectrum was then square-rooted and averaged across the frequency band of 0.01–0.08 Hz at each voxel, which represents ALFF. Finally, the amplitudes (beta scores) of subject-level maps were transformed into Z-scores to create standardized subject-level maps for each participant in the EO and in the EC condition for the statistical analysis.

### Data analysis

To determine whether we were able to replicate previous results assessed with ALFF in the EO compared to the EC condition in typically sighted individuals^35-37,66^ a voxel-wise paired t-test was carried out in the group of 28 typically sighted individuals comparing the eyes open and the eyes closed condition (Fig. 1; Supplementary Table 1). The statistical map was corrected for multiple comparisons using a Gaussian Random Field (GRF) correction (voxel-wise *p* < .01, cluster-wise *p* < .05, corrected).^67^

**Figure 1.**
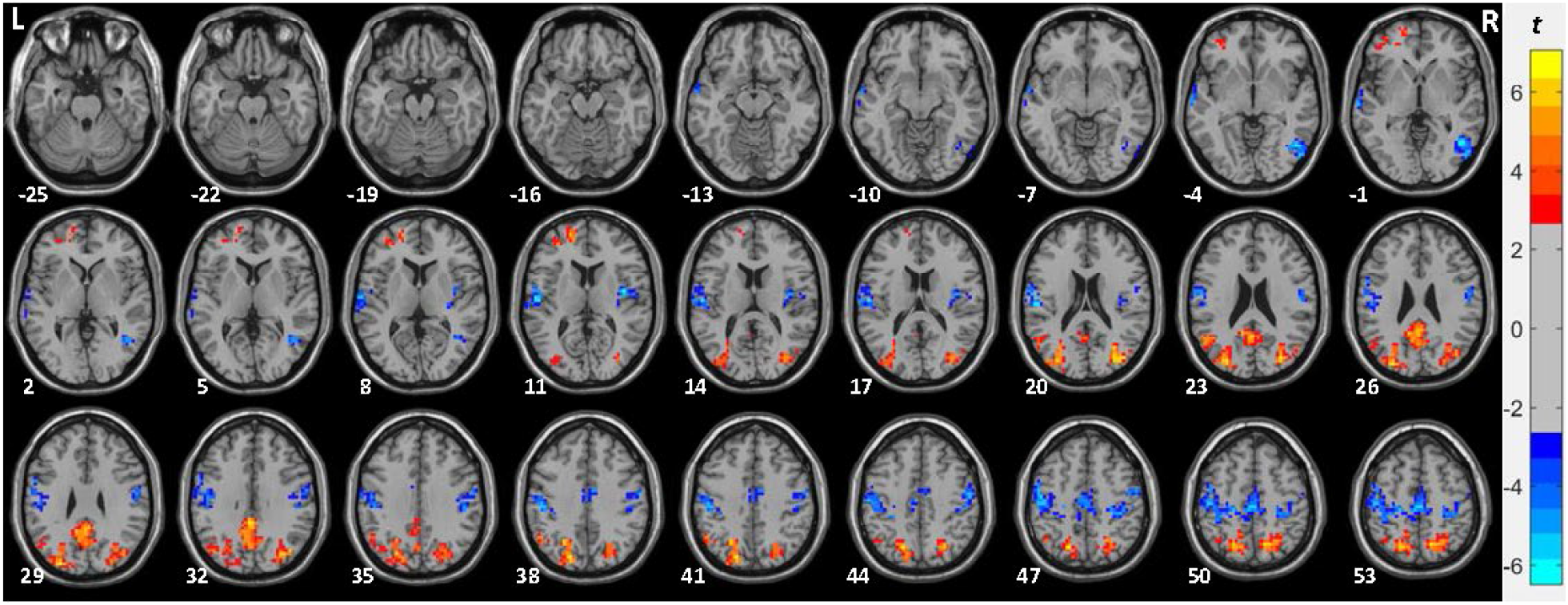
ALFF in the EO compared to the EC condition in sighted control individuals (SC group, n=28). Paired t-test results of the amplitude of low frequency fluctuations (ALFF) comparing the eyes open (EO) and the eyes closed (EC) condition in the group of sighted control individuals (SC group, n=28). The red colors denote voxels with significantly higher amplitude in the EO compared to the EC condition and the blue colors denote voxels with significantly lower amplitude in the EO compared to the EC condition. Significant clusters are shown after Gaussian random field (GRF) correction for multiple spatial comparisons (voxel-wise *p* < .01, cluster-wise *p* < .05, corrected).

To determine the brain regions with significantly higher and lower ALFF between the EO compared to the EC condition in each of the tested groups, voxel-wise paired t-tests were separately calculated for: 1) the SC group matched in sex and age to the CC group, 2) the CC group, 3) the CB group and 4) the DC group (Fig. 2; Supplementary Fig. 3A-C; Supplementary Table 2). The statistical maps were corrected for multiple comparisons using a Gaussian Random Field (GRF) correction (voxel-wise *p* < .01, cluster-wise *p* < .05, corrected).^67^

**Figure 2.**
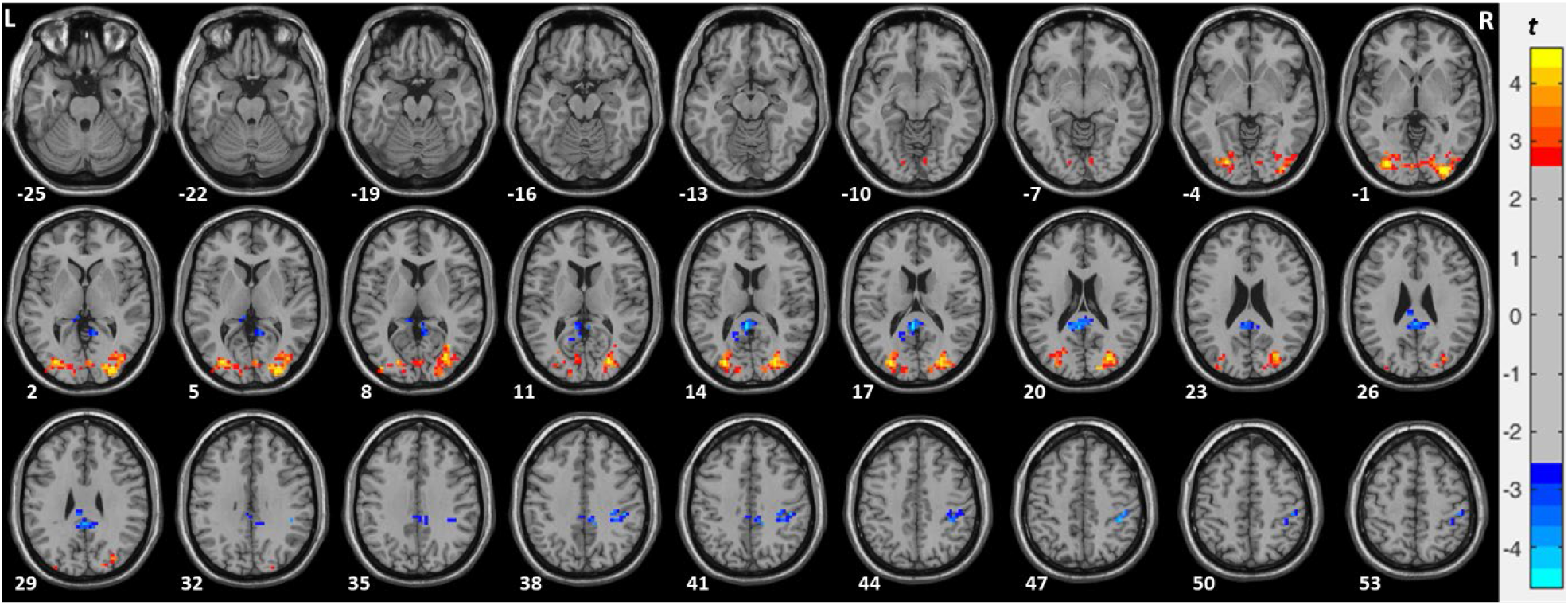
ALFF in the EO compared to the EC condition in CC individuals (CC group, n=19). Paired t-test results of the amplitude of low frequency fluctuations (ALFF) comparing the eyes open (EO) and the eyes closed (EC) condition in the group of congenital cataract-reversal individuals (CC group, n=19). The red colors denote voxels with significantly higher amplitude in the EO compared to the EC condition and the blue colors denote voxels with significantly lower amplitude in the EO compared to the EC condition. Significant clusters are shown after Gaussian random field (GRF) correction for multiple spatial comparisons (voxel-wise *p* < .01, cluster-wise *p* < .05, corrected).

To assess group differences in ALFF between the eyes open compared to the eyes closed condition, a 2 x 2 mixed effect model on standardized ALFF maps (Z-scores) was carried out. Three group models were run: (1) CC group vs SC group; (2) DC group vs. SC group; (3) CB group vs. SC group (Fig. 3; Table 4; Supplementary Fig. 1A-B). Note that in a mixed effect model, we matched typically sighted individuals approximately in age and sex to the participants of each tested group (see *Participants* section). All interaction maps were corrected for multiple comparisons using the GRF-correction with a threshold of voxel-wise *p* < .05, cluster-wise *p* < .025, corrected.^68-70^

**Figure 3.**
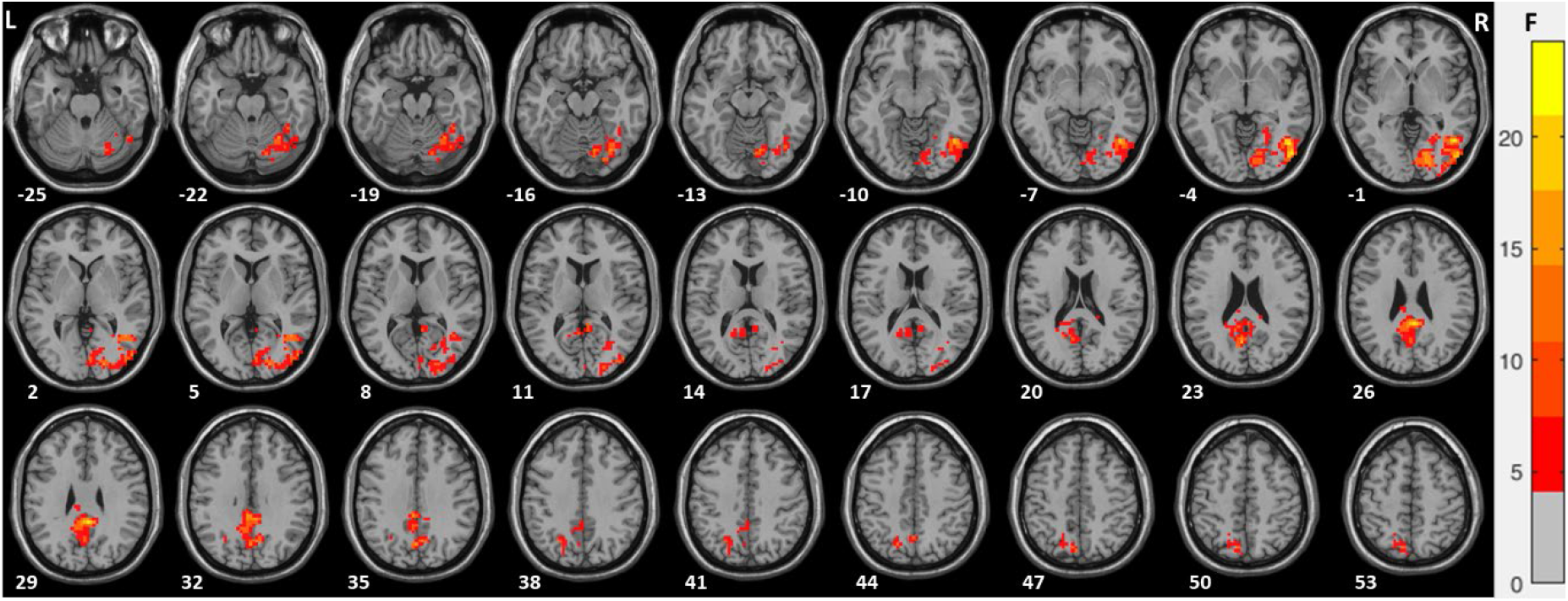
Group differences in ALFF in the EO compared to the EC condition in CC individuals compared to the SC group (n=19). Using standardized ALFF (amplitude of low frequency fluctuations) i.e. Z-scores, a mixed 2×2 model (group x condition) was carried out for congenital cataract-reversal individuals (CC) vs. the SC group (n=19). Regions with significant interaction effects are shown after Gaussian random field theory (GRF) correction for multiple spatial comparisons (voxel-wise *p* < .05, cluster-wise *p* < .025, corrected).

**Table 4.**
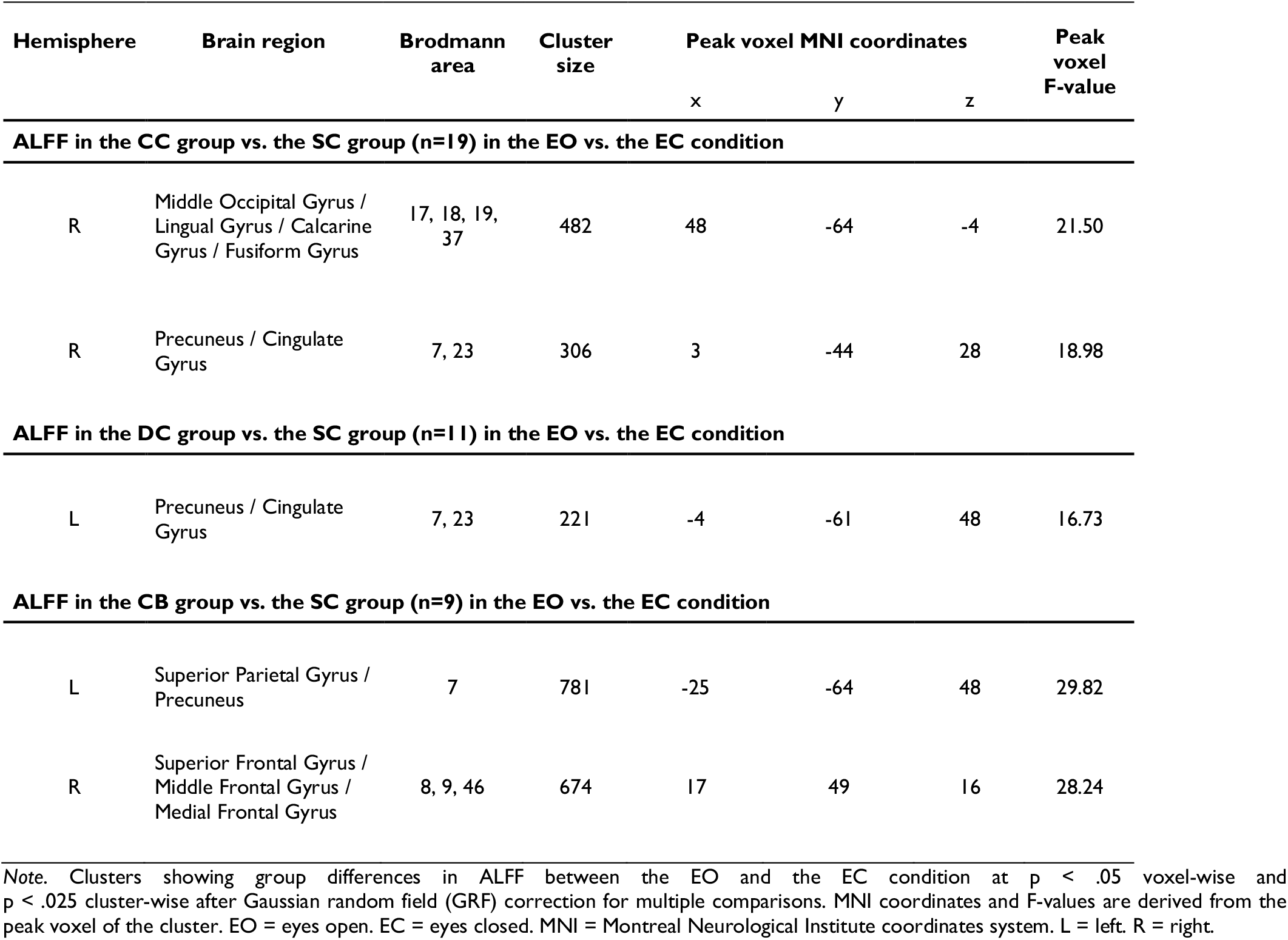
Group differences in ALFF in the EO compared to the EC condition.

| Hemisphere | Brain region | Brodmann area | Cluster size | Peak voxel MNI coordinates |  |  | Peak voxel F-value |
| --- | --- | --- | --- | --- | --- | --- | --- |
|  |  |  |  | x | y | z |  |
| ALFF in the CC group vs. the SC group (n=19) in the EO vs. the EC condition |  |  |  |  |  |  |  |
| R | Middle Occipital Gyrus / Lingual Gyrus / Calcarine Gyrus / Fusiform Gyrus | 17, 18, 19, 37 | 482 | 48 | -64 | -4 | 21.50 |
| R | Precuneus / Cingulate Gyrus | 7, 23 | 306 | 3 | -44 | 28 | 18.98 |
| ALFF in the DC group vs. the SC group (n=11) in the EO vs. the EC condition |  |  |  |  |  |  |  |
| L | Precuneus / Cingulate Gyrus | 7, 23 | 221 | -4 | -61 | 48 | 16.73 |
| ALFF in the CB group vs. the SC group (n=9) in the EO vs. the EC condition |  |  |  |  |  |  |  |
| L | Superior Parietal Gyrus / Precuneus | 7 | 781 | -25 | -64 | 48 | 29.82 |
| R | Superior Frontal Gyrus / Middle Frontal Gyrus / Medial Frontal Gyrus | 8, 9, 46 | 674 | 17 | 49 | 16 | 28.24 |

Note. Clusters showing group differences in ALFF between the EO and the EC condition at $p < .05$ voxel-wise and $p < .025$ cluster-wise after Gaussian random field (GRF) correction for multiple comparisons. MNI coordinates and F-values are derived from the peak voxel of the cluster. EO = eyes open. EC = eyes closed. MNI = Montreal Neurological Institute coordinates system. L = left. R = right.

Since eyes open vs. closed does not mean the same for the CB group as for the other groups, and to identify whether the CC group shows different resting state activity particularly in the EO condition (see hypotheses), the CC group and their matched SC group were separately compared in the EO and in the EC condition using voxel-wise t-tests (Fig. 4A; Supplementary Table 3). The analogous analyses were run for the CB group and the DC group (Fig. 4B and Supplementary Fig. 2A-B; Supplementary Table 3). Additionally, the CC group and the CB group were separately compared in the EO and in the EC condition using voxel wise t-tests (Supplementary Fig. 4; Supplementary Table 4). The statistical maps were corrected for multiple comparisons using a Gaussian Random Field (GRF) correction (voxel-wise *p* < .01, cluster-wise *p* < .05, corrected).^67^

**Figure 4.**
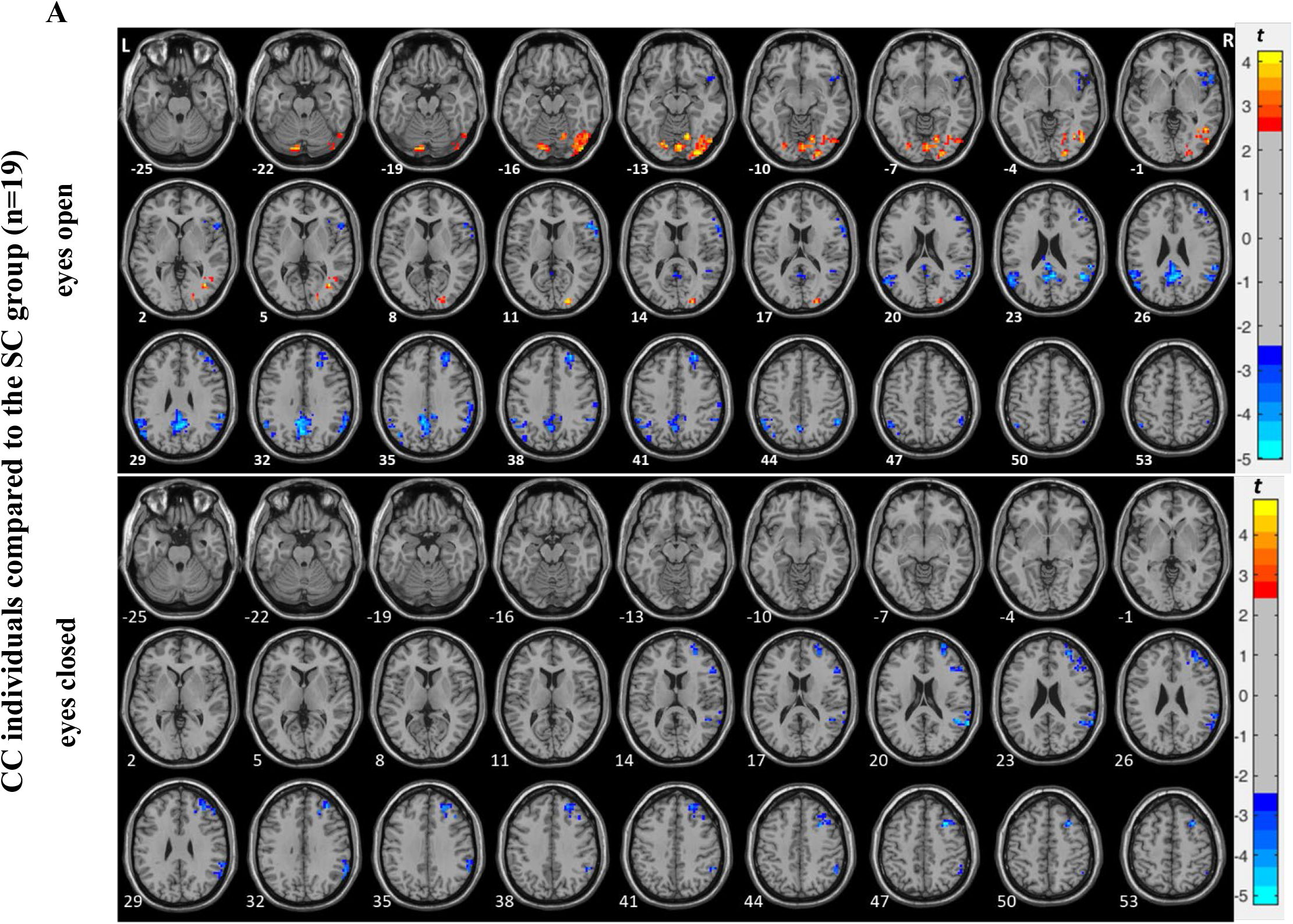
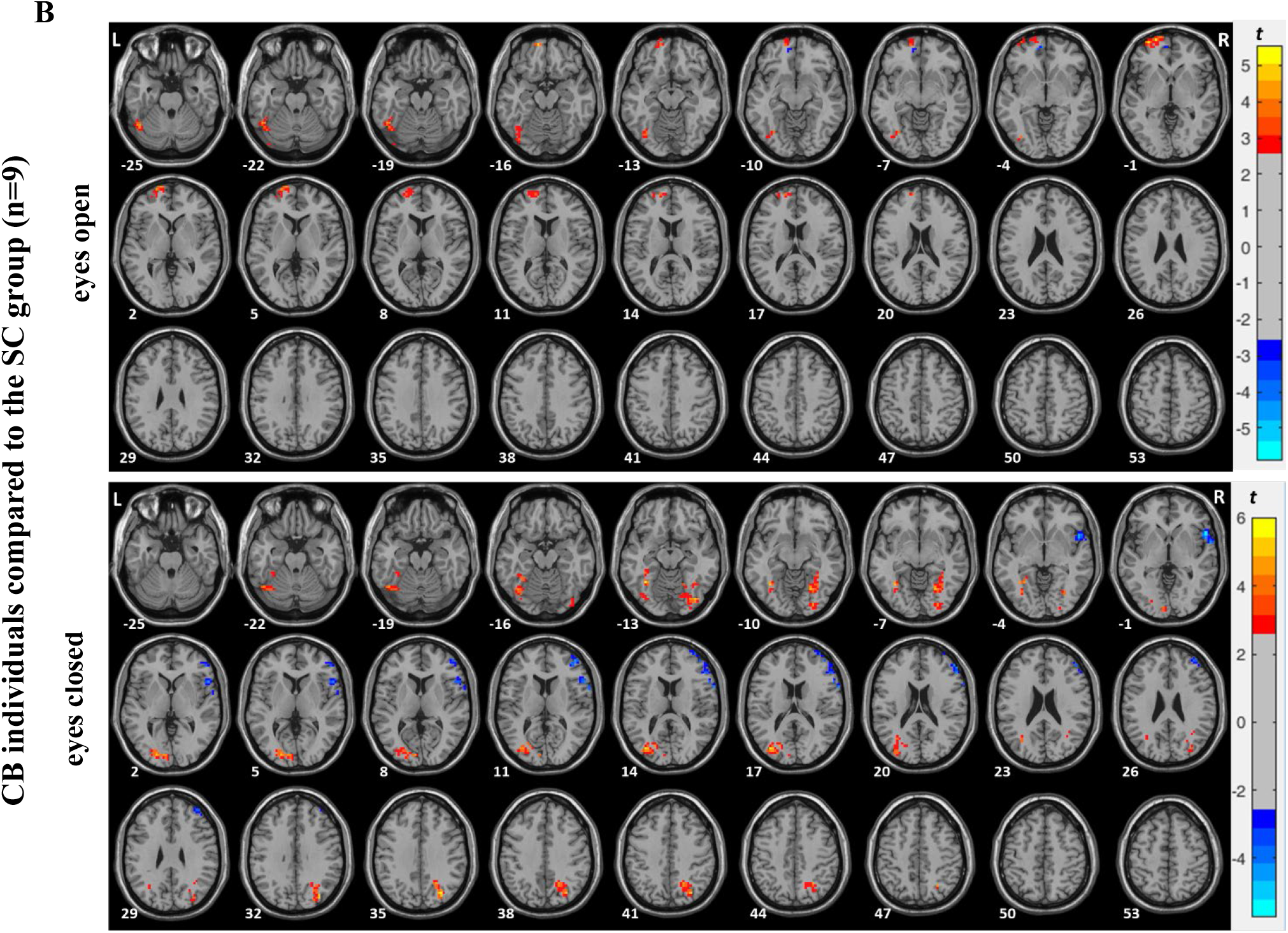
ALFF in the EO and in the EC condition. Two-sample t-test results of the amplitude of low frequency fluctuations (ALFF) comparing **(A)** congenital cataract-reversal individuals (CC) and the SC group (n=19) matched in age and sex to the CC group and **(B)** congenitally blind individuals (CB) and the SC group (n=9) matched in age and sex to the CB group in the **eyes open (EO)** condition and in the **eyes closed (EC)** condition. The red colors denote voxels with significantly higher amplitude for (A) the CC compared to the SC group and for (B) the CB compared to the SC group and the blue colors denote voxels with significantly lower amplitude for (A) the CC compared to the SC group and for (B) the CB compared to the SC group. Significant clusters are shown after Gaussian random field theory (GRF) correction for multiple spatial comparisons (voxel-wise *p* < .01, cluster-wise *p* < .05, corrected).

A probabilistic cytoarchitectonic atlas of the human brain (Automated Anatomical Labeling, Harvard-Oxford Cortical and Subcortical Structural Atlases, and Brodmann Atlas) as implemented in the DPABI toolbox, was used to assign significant voxels to brain regions.

### Data availability

Anonymized data and materials will be made available to the external scientists upon reasonable request to the corresponding author through data transfer agreements approved by the stakeholders, under stipulations of applicable law including but not limited to the General Data Protection Regulation (EU 2016/679).

## Results

### ALFF in the EO compared to the EC condition in sighted control individuals (SC group, n=28)

To determine brain regions with significantly higher and lower ALFF in the EO compared to the EC condition in sighted controls voxel-wise paired t-test was carried out (with correction for multiple comparisons see *Data analysis* section). We found significantly higher ALFF in a cluster in left visual association areas (BA7, 19), as well as in a cluster in the right visual association areas (BA7, 19) and in a cluster in the left precuneus (BA7). Additionally, ALFF was higher in the EO than in the EC condition in a cluster in the left frontal cortex (BA8, 9, 46). ALFF was significantly lower in the EO than in the EC condition bilaterally in sensorimotor and temporal (auditory) regions i.e. in clusters covering pre (BA4, 6) and postcentral gyrus (BA3), middle temporal gyrus, superior temporal gyrus and inferior temporal gyrus (BA20, 21, 22). Additionally, ALFF was significantly lower in the EO than in the EC condition in a cluster in the left frontal regions (BA4, 8, 9) (see Supplementary Table 1 and Fig. 1 for more details).

These results, by and large, replicated previously reported findings.^35-37,66^

### ALFF in the CC group vs. the SC group (n=19) in the EO vs. the EC condition

To determine brain regions with significantly higher and lower ALFF in <u>congenital cataract-reversal individuals</u> in the EO compared to the EC condition voxel-wise paired t-test was carried out. We found significantly higher ALFF in the EO compared to the EC condition in a cluster in the right and left visual cortex including the calcarine gyrus (BA17), lingual gyrus (BA18) and the middle occipital gyrus (BA19). Significantly lower ALFF in the EO compared to the EC condition was found in a cluster in the left cingulate gyrus (BA23) and in a cluster in the supramarginal (BA40) and postcentral gyrus (BA2, 3) (see Fig. 2 and Supplementary Table 2).

To test for possible group differences, the interaction of group and condition was analyzed using a 2 x 2 mixed effect model (CC group vs. SC group x EO condition vs. EC condition) with the standardized ALFF maps (Z-scores) as the dependent variable. A significant interaction was found in a large cluster in right visual areas spanning the calcarine gyrus (BA17), lingual gyrus (BA18), middle occipital gyrus (BA19), and the fusiform gyrus (BA37) and in a cluster predominantly in the precuneus (BA7) (see Fig. 3 and Table 4 for more details).

We next compared the CC and the SC group separately in the EO and in the EC condition with voxel-wise two-sample t-tests. In the <u>eyes open condition</u>, we observed significantly higher ALFF in the CC group than in the SC group in a cluster in the right visual cortex including the calcarine gyrus (BA17), lingual gyrus (BA18) and the inferior and middle occipital gyrus (BA19). Significantly lower ALFF in the CC group than in the SC group was observed in a cluster in the right precuneus (BA7), and in clusters including parietal (BA39, 40), temporal (BA22), and frontal regions (BA8, 9, 45, 46) (see Supplementary Table 3; Fig. 4A upper panel: eyes open). In the <u>eyes closed condition</u> we did not observe any region with significantly higher ALFF in the CC group than in the SC group. Significantly lower ALFF in the CC group than in the SC group was observed in the right hemisphere in a cluster in frontal regions (BA8, 9, 46) and in a cluster in temporal (BA22) and parietal regions (BA39, 40) (see Supplementary Table 3 and Fig. 4A lower panel: eyes closed).

### ALFF in the DC group vs. the SC group (n=11) in the EO vs. the EC condition

To determine brain regions with significantly higher and lower ALFF in <u>developmental cataract-reversal individuals</u> in the EO compared to the EC condition voxel-wise paired t-test was carried out. Significantly higher ALFF in the EO compared to the EC condition in the DC group was observed bilaterally in several clusters of the frontal cortex (BA8, 9, 45, 46). Significantly lower ALFF in the EO compared to the EC condition was observed in a cluster in the right early visual areas i.e. in the calcarine gyrus (BA17) and lingual gyrus (BA18), and in a cluster in temporal regions (BA21). Additionally, lower ALFF was found in a cluster in sensorimotor areas including the pre (BA4) and postcentral gyrus (BA2, 3) (Supplementary Fig. 3B; Supplementary Table 2).

A 2 x 2 mixed effect model (DC group vs. SC group x EO condition vs. EC condition) was calculated with standardized ALFF maps (Z-scores) as the dependent variable to test the interaction between group and condition. A significant interaction was found in a cluster in parietal cortex (BA7) (Supplementary Fig. 1A; Table 4).

We next compared ALFF between the DC and the SC group separately in the EO and in the EC condition with voxel-wise two-sample t-tests. In the <u>eyes open condition</u>, we observed significantly lower ALFF in the DC group than in the SC group in a cluster in parietal cortex, predominantly in the precuneus (BA7). We did not observe any region with significantly higher ALFF in the DC group compared to the SC group (Supplementary Table 3; Supplementary Fig. 2A). In the <u>eyes closed condition</u> we did not observe any region with significantly higher or lower ALFF in the DC compared to the SC group (Supplementary Fig. 2B).

### ALFF in the CB group vs. the SC group (n=9) in the EO vs. the EC condition

To determine brain regions with significantly higher and lower ALFF in <u>congenitally blind individuals</u> in the EO compared to the EC condition voxel-wise paired t-test was carried out. We found significantly higher ALFF in the EO compared to the EC condition in a cluster in frontal regions (BA8, 9, 46). Significantly lower ALFF in the EO compared to the EC condition was found in a cluster in parietal regions (BA7) (Supplementary Fig. 3C; Supplementary Table 2).

To analyze the interaction of group and condition a 2 x 2 mixed effect model (CB group vs. SC group x EO condition vs. EC condition) was carried out with standardized ALFF (Z-scores) as the dependent variable. Significant interaction effects were observed in a cluster in parietal regions (BA7) and in a cluster in frontal regions (BA8, 9, 46) (Supplementary Fig. 1B; Table 4).

We then compared the CB group and SC group separately in the EO and in the EC condition with voxel-wise two-sample t-tests. In the <u>eyes open condition</u>, we found significantly higher ALFF in the CB group than in the SC group in a cluster in the left visual areas spanning middle occipital gyrus (BA19) and fusiform gyrus (BA37) and in a cluster in frontal regions (BA8, 9).

We did not observe any regions with significantly lower ALFF for the CB group compared to the SC group (Supplementary Table 3; Fig. 4B upper panel: eyes open). In the <u>eyes closed condition</u> we observed significantly higher ALFF in the CB group than in the SC group in two clusters in the left visual cortex: one in the middle occipital gyrus (BA19) and one in the fusiform gyrus (BA37) and in two clusters in right visual areas: one in dorsal visual association cortex (BA7, 19) and one in ventral visual cortex (BA18, 37). Significantly lower ALFF in the CB group than in the SC group was found in the right hemisphere in two clusters: one in frontal regions (BA45, 46) and one in frontal-temporal regions (BA22, 45) (Supplementary Table 3; Fig. 4B lower panel: eyes closed).

We next (exploratively) compared the CC and the CB group separately in the EO and in the EC condition with voxel-wise two-sample t-tests. In the <u>eyes open condition</u>, we observed significantly higher ALFF in the CC group than in the CB group in a cluster in the superior frontal gyrus (BA8, 9). Significantly lower ALFF in the CC group than in the CB group was found in two clusters in parietal cortex (BA39, 40) (Supplementary Table 4; Supplementary Fig. 4A). In the <u>eyes closed condition</u>, we observed significantly lower ALFF in the CC group than in the CB group in the right hemisphere in a cluster located in early visual cortex and visual association areas spanning calcarine gyrus (BA17), lingual gyrus (BA18), middle occipital gyrus (BA19) and fusiform gyrus (BA37), and in the left hemisphere in two clusters: one in visual association areas spanning middle occipital gyrus (BA19), fusiform gyrus (BA37) and including parts of the cerebellum and one located in early visual areas and visual association areas spanning calcarine gyrus (BA17), lingual gyrus (BA18) and middle occipital gyrus (BA19). We did not observe any regions with significantly higher ALFF in the CC group than in the CB group (Supplementary Table 4; Supplementary Fig. 4B).

## Discussion

The goal of the present study was to identify whether the emergence of typical resting state activity of the human brain, as the prerequisite of any task related processing, depends on experience during a sensitive period of early brain development. To this end, we tested to which degree BOLD resting state activity in an eyes open (EO) and eyes closed (EC) condition recovered after a transient phase of congenital blindness due to congenital cataracts. Congenital cataract-reversal individuals (CC group) were compared to normally sighted controls (SC group) and to a group of congenitally blind humans (CB group). Developmental cataract-reversal individuals (DC group) served as additional control group. All groups were investigated in the same scanner, came from the same community and the groups were matched in age.

First, we replicated the typical ALFF (Amplitude of Low Frequency Fluctuations) pattern in the SC group^35-37,66^: ALFF was significantly higher in the EO than in the EC condition mostly in visual association cortex and in parietal cortex. Moreover, ALFF was significantly lower in the EO than in the EC condition in sensorimotor and auditory cortices.

Importantly, ALFF varied with EO vs. EC in the CC group’s visual cortices as well: Similar to the SC group the amplitude of slow BOLD fluctuations was higher in the EO than in the EC condition. However, there were several group differences too: In the EO condition visual cortex activity was overall higher in the CC than in the SC group. Moreover, in the CC group an increase of ALFF in parietal cortex was not observed in the EO compared to the EC condition. In fact ALFF was lower in the CC group than in the SC group in the EO condition. Moreover, the decrease of ALFF for the EO compared to the EC condition in auditory and sensorimotor regions was missing in the CC group. Finally, the CB group showed higher ALFF than SC individuals in both the EO and the EC condition, and higher ALFF than the CC group in the EC condition.

Research in non-human primates showed that synaptic pruning in visual cortex is experience dependent and particularly affects the asymmetric, excitatory synapses, resulting in an experience dependent set-point for visual cortex excitability. Cortical thickness development runs parallel to the developmental trajectory of synaptogenesis.^71^ In fact, permanently congenitally blind individuals feature thicker visual cortices which was interpreted as indicating an arrest of experience dependent synaptic pruning.^72-76^ Importantly, a higher cortical thickness has recently been observed in CC individuals^77,78^ too. These results thus suggest that the process of synaptic pruning in early visual cortex is linked to a sensitive period in early primate brain development. The presence of exuberant synapses has been demonstrated to result in higher glucose uptake^7,79^ and presumably blood flow during rest.^80^ Thus, we speculate that higher ALFF might reflect higher resting state excitatory activity of less pruned neural circuits within the occipital lobe of CC (and CB) individuals.

Early visual cortex in sighted individuals is characterized by a high degree of inhibition, which results in a short time constant and thus the ability to process visual information at a fast rate.^81^ Non-human animal research has demonstrated that the elaboration of inhibitory neural networks is a hallmark of sensitive period plasticity.^82^ In fact, stabilization of inhibitory synapse and myelination ends the sensitive period.^83^

Previous EEG studies in CB and CC individuals have repeatedly observed lower alpha oscillatory activity^51,53,54^ and in CB individuals higher gamma oscillatory activity.^55,56^ Alpha oscillatory activity has been considered to be an electrophysiological signature for the control of the excitatory/inhibitory (E/I) balance of neural circuits.^84^ In the present context, it is important to note that alpha oscillatory activity has been found to inversely correlate with ALFF,^45-47,85^ while gamma band activity was found in monkeys to positively correlated with slow BOLD fluctuations.^59^ Thus, the higher posterior ALFF observed for the CC and the CB groups in the present study is consistent with reduced posterior alpha band activity in these groups and higher gamma band activity reported for CB individuals. All these findings converge to the hypothesis that overall excitation is enhanced in the visual cortex of CB and CC individuals. Moreover, the visually triggered BOLD signal seems to be correlated with changes in the glutamate level.^86^ In fact, there is evidence of higher glutamatergic^87^ and lower GABAergic activity in congenitally permanently blind individuals.^88^ Corresponding data in CC individuals are not available yet.

However, there was a crucial difference between the CC and the CB group. Higher ALFF was observed for the CC group than in the SC group only in the EO condition and in fact in the EC condition ALFF was lower in the CC than in the CB group. These group differences demonstrate that the visual cortical networks partially recovered in the CC group, that is, different resting state activity levels were adopted as a function of whether or not light reaches the retina during rest.

Visual thalamo-cortical input excites pyramidal neurons in the granular layers of the cortex but in parallel entertains synapses to inhibitory interneurons, which allows a quick shutting down of excitation. From non-human animal research, it is known that these inhibitory circuits are shaped by experience and that they are stabilized by perineural networks^82^ resulting in neural circuits^89^ which selectively respond only to certain input. Thus, we speculate that higher ALFF in the CC group in the EO condition indicates less selective processing and an impaired quick shutting down of visual driven activation possibly due to a compromised intracortical inhibition. In fact, behavioral studies have shown longer lasting visual (motion) aftereffects in CC individuals,^19^ which indirectly supports this speculation. By contrast, we hypothesize that higher ALFF in visual cortex in the CB group was predominantly due to higher spontaneous activity. Higher spontaneous activity in visual areas is an often reported finding in visually deprived non-human animals.^90,91^ In fact, a study in monkeys has observed a decrease of spontaneous activity in visual cortex after ending a phase of congenital lid suture.^90^ Thus, a decrease in spontaneous activity might explain the lower ALFF in visual areas in the CC than in the CB group during the EC condition and might explain as well indistinguishable ALFF of the CC and the SC groups in the EC condition. Lower spontaneous activity would be compatible with the idea of partial E/I balance recovery in visual cortex.

In sum, we suggest that higher slow BOLD fluctuations in CB and CC individuals might originate from a similar neural substrate, that is, not or less well attuned visual circuits. However, despite late availability of patterned visual input, the neural circuits in the CC group seem to had recovered to some degree too, such that spontaneous activity decreased. However, the fine-tuned neural (inhibitory) circuits which allow for a selective activation and quick shutting down of stimulus-driven activity might not have fully emerged, resulting in an enhanced and possibly longer-lasting excitation as a response to visual stimulation.

Importantly, we interpret our results on slow BOLD fluctuations in the visual cortices of CC individuals as evidence for retracted cross-modal plasticity. In fact, in the context of cross-modal plasticity in deaf cats it has been argued that the higher level of excitation in auditory cortex reflects a largely reduced threshold in order to allow for cross-modal activation.^92^ Thus, visual entrainment of visual areas in the CC group might have enhanced the threshold for cross-modal activation and reduced spontaneous activity possibly via homeostatic plasticity mechanisms.^93^

In parietal cortex we observed a lower ALFF in the CC than in the SC group in the EO condition. Hyvärinen et al.^94^ reported a lower visual responsiveness in parietal area BA7 in monkeys who had been visually deprived for 7 to 11 months. In a follow-up study one year after the end of the deprivation period responsiveness to visual stimulation had further declined rather than increased, as would have been expected from restoring sight.^95^ Except one participant (assessed 6 months post cataract removal surgery) all CC participants of the present study were assessed more than one year after cataract removal surgery. Here, we speculate that the observed lower resting state activity in the CC individuals in parietal regions during EO might reflect a similar reduced regain of visual responsiveness in parietal cortex as observed by Hyvärinen et al.^94^ in non-human primates. Although parietal cortex is a multisensory region, many processes including multisensory spatial integration are visually dominated in sighted individuals. Thus, lower ALFF in parietal cortex in the CC compared to the SC group might indicate a lower visual influence on multisensory (spatial) processing. In fact, two behavioral studies have found a reduced visual impact on tactile spatial performance in congenital cataract-reversal individuals with a history of longer lasting visual deprivation.^96,97^

Altered multisensory processing is suggested by the third main result in the CC group too: Activity in auditory and sensorimotor regions was, in contrast to the SC group (and the DC group) not lower during EO than during EC (see Fig. 2). Functional connectivity studies in sighted humans have provided ample evidence for a higher functional coupling of visual and auditory as well as visual and sensorimotor cortices during EC than during EO.^98^ Crucially, such overall coupling between visual brain regions and both auditory and sensorimotor brain regions seems to be reduced in congenitally blind humans.^62^ Our new finding that resting state activity in auditory and sensorimotor regions is unaffected by eyes opening in CC individuals might suggest, analogously to parietal cortex, a reduced impact of vision in multisensory processing. This idea is compatible with the previously reported lower lip reading specific activity in the superior temporal sulcus of CC individuals^99^ and the lack of audio-visual enhancements neither in this region^15^ nor in behavior.^100^

Finally, it has to be noted that similar group differences as found between the CC and the SC group were not observed for the DC group except the lower parietal cortex activity during EO. By contrast, the typical decrease in ALFF for resting state activity with EO vs. EC was highly robust in the DC group. Thus, a typical modulation of auditory and sensorimotor cortex activity by the visual system might crucially depend on connectivity elaborated in early brain development. By contrast, the lower impact of vision on parietal cortex activity condition might reflect a lower online weighting due to overall reduced reliability of visual input.^101^

## CONCLUSION

Slow BOLD fluctuations indicating resting state activity of neural circuits suggested a retraction of cross-modal plasticity after sight restoration in individuals with a history of blindness due to congenital cataracts. However, visual neural circuits seemed to be less tuned and activity thus was not as well-regulated as in normally sighted controls. This impairment might reflect remaining visual cortical circuit changes as a consequence of congenital blindness.

A significant influence of the visual system on parietal as well as auditory and sensorimotor systems as typically found in sighted individuals had not recovered in the congenital cataract-reversal individuals, suggesting a high influence of experience on multisensory neural networks.

Since resting state brain activity builds the scaffold for task related processing, we put forward the hypothesis that the incomplete recovery of typical resting state activity patterns within the visual system and across sensory systems might contribute to the persisting visual and multisensory impairments after restoring sight in people with a congenital loss of pattern vision.

## Acknowledgments

We thank, Rakesh Balanchadar, and Seema Banerjee, Divya Jagadish for help with participant recruitment and data acquisition, as well as Armin Heinecke for an initial data quality assessment. We are grateful to Kabilan Pitchaimuthu, Prativa Regmi and Idris Shareef for providing clinical details of the participants during the classification process and Suddha Sourav for technical support. We thank the technical staff of the Lucid Medical Diagnostics Banjara Hills in Hyderabad in India, in particular Balakrishna Vaddepally, for technical assistance during collection of MRI data. We are grateful to D. Balasubramanian of the L.V. Prasad Eye Institute for initiating and supporting our research.

## Supplementary material

Supplementary material is available online.

## Abbreviations

BA: Brodmann area
CB: congenitally blind individuals
CC: congenital cataract-reversal individuals
DC: developmental cataract-reversal individuals
DICOM: Digital Imaging and Communications in Medicine
EC: eyes closed
EO: eyes open
EPI: echo-planar imaging
FA: flip angle
fMRI: functional magnetic resonance imaging
FOV: Field Of View
FWHM: full width at half maximum
logMAR: Logarithm of the Minimum Angle of Resolution
MEG: Magnetoencephalography
NIFTI: Neuroimaging Informatics Technology Initiative
SC: sighted control individuals
TE: echo time
TR: repetition time

## References

1. Röder B, Neville HJ. Developmental plasticity. In Grafman J, Robertson IH, eds. Handbook of neuropsychology, 2nd revised edition. else; 2003;9:231–270.

2. Röder B, Kekunnaya R, Guerreiro MJ. Neural mechanisms of visual sensitive periods in humans. Neuroscience and Biobehavioral Reviews. 2021;120:86–99.

3. Elbert T, Sterr A, Rockstroh B, Pantev C, Müller MM, Taub E. Expansion of the tonotopic area in the auditory cortex of the blind. Journal of Neuroscience. 2002;22(22):9941–9944.

4. Niemeyer W, Starlinger I. Do the blind hear better? Investigations on auditory processing in congenital or early acquired blindness II. Central functions. Audiology. 1981;20(6):510–515.

5. Röder B, Rösler F, Neville HJ. Effects of interstimulus interval on auditory event-related potentials in congenitally blind and normally sighted humans. Neuroscience Letters. 1999;264(1-3):53-56.

6. Veraart C, De Volder AG, Wanet-Defalque MC, Bol A, Michel C, Goffinet AM. Glucose utilization in human visual cortex is abnormally elevated in blindness of early onset but decreased in blindness of late onset. Brain Research. 1990;510(1):115–121.

7. Wanet-Defalque MC, Veraart C. De Volder A, et al. High metabolic activity in the visual cortex of early blind human subjects. Brain Research. 1988;446(2):369–373.

8. Bedny M. Evidence from blindness for a cognitively pluripotent cortex. Trends in Cognitive Sciences. 2017;21(9):637–648.

9. Renier L, De Volder AG, Rauschecker JP. Cortical plasticity and preserved function in early blindness. Neuroscience & Biobehavioral Reviews. 2014;41:53–63.

10. Heimler B, Amedi A. Are critical periods reversible in the adult brain? Insights on cortical specializations based on sensory deprivation studies. Neuroscience & Biobehavioral Reviews. 2020;116: 494–507.

11. Merabet LB, Pascual-Leone A. Neural reorganization following sensory loss: the opportunity of change. Nature Reviews Neuroscience. 2010;11(1):44–52.

12. Voss P, Collignon O, Lassonde M, Lepore F. Adaptation to sensory loss. Wiley Interdisciplinary Reviews: Cognitive Science. 2010;1(3):308–328.

13. Sinha P., Held R. Sight restoration. F1000 Medicine Reports. 2012;4:17.

14. Bottari D, Kekunnaya R, Hense M, Troje NF, Sourav S, Röder B. Motion processing after sight restoration: No competition between visual recovery and auditory compensation. NeuroImage. 2018;167:284–296.

15. Guerreiro MJ, Putzar L, Röder B. The effect of early visual deprivation on the neural bases of multisensory processing. Brain. 2015;138(6):1499–1504.

16. Collignon O, Dormal G, De Heering A, Lepore F, Lewis TL, Maurer D. Long-lasting crossmodal cortical reorganization triggered by brief postnatal visual deprivation. Current Biology. 2015;25(18):2379–2383.

17. Saenz M, Lewis LB, Huth AG, Fine I, Koch C. Visual motion area MT+/V5 responds to auditory motion in human sight-recovery subjects. Journal of Neuroscience. 2008;28(20):5141–5148.

18. Huber E, Chang K, Alvarez I, Hundle A, Bridge H, Fine I. Early blindness shapes cortical representations of auditory frequency within auditory cortex. Journal of Neuroscience. 2019;39(26):5143–5152.

19. Guerreiro MJ, Putzar L, Röder B. Persisting cross-modal changes in sight-recovery individuals modulate visual perception. Current Biology. 2016;26(22):3096–3100.

20. Kral A, Land R, Baumhoff P, Tillein J, Hubka P, Lomber SG. Cross-modal plasticity in the congenitally deaf cat. Multisensory Research. 2013;26:37–37.

21. Land R, Baumhoff P, Tillein J, Lomber SG, Hubka P, Kral A. Cross-modal plasticity in higher-order auditory cortex of congenitally deaf cats does not limit auditory responsiveness to cochlear implants. Journal of Neuroscience. 2016;36(23):6175–6185.

22. Collignon O, Vandewalle G, Voss P, et al. Functional specialization for auditory–spatial processing in the occipital cortex of congenitally blind humans. Proceedings of the National Academy of Sciences. 2011;108(11):4435–4440.

23. Röder B, Kekunnaya R. Visual experience dependent plasticity in humans. Current Opinion in Neurobiology. 2021;67:155–162.

24. Ganesh S, Arora P, Sethi S, et al. Results of late surgical intervention in children with early-onset bilateral cataracts. British Journal of Ophthalmology. 2014;98(10):1424–1428.

25. Levelt CN, Hübener M. Critical-period plasticity in the visual cortex. Annual Review of Neuroscience. 2012;35:309–330.

26. Hensch, TK. Critical Period Regulation. Annual Review of Neuroscience. 2004;27:549–579.

27. Berkes P, Orbán G, Lengyel M, Fiser J. Spontaneous cortical activity reveals hallmarks of an optimal internal model of the environment. Science 2011;331(6013):83–87.

28. Biswal B, Zerrin Yetkin F, Haughton VM, Hyde JS. Functional connectivity in the motor cortex of resting human brain using echo-planar MRI. Magnetic Resonance in Medicine. 1995;34(4):537–541.

29. Fox MD, Snyder AZ, Vincent JL, Corbetta M, Van Essen DC, Raichle ME. The human brain is intrinsically organized into dynamic anticorrelated functional networks. Proceedings of the National Academy of Sciences. 2005;102(27):9673–9678.

30. Fox MD, Raichle ME. Spontaneous fluctuations in brain activity observed with functional magnetic resonance imaging. Nature Reviews Neuroscience. 2007;8(9):700–711.

31. Deco G, Corbetta M. The dynamical balance of the brain at rest. The Neuroscientist. 2011;17(1):107–123.

32. Honey CJ, Sporns O, Cammoun L, et al. Predicting human resting-state functional connectivity from structural connectivity. Proceedings of the National Academy of Sciences. 2009;106(6): 2035–2040.

33. Vincent JL, Patel GH, Fox MD. Intrinsic functional architecture in the anaesthetized monkey brain. Nature. 2007;447(7140):83–86.

34. Yu-Feng Z, Yong H, Chao-Zhe Z, et al. Altered baseline brain activity in children with ADHD revealed by resting-state functional MRI. Brain and Development. 2007;29(2):83–91.

35. Yang H, Long XY, Yang Y, et al. Amplitude of low frequency fluctuation within visual areas revealed by resting-state functional MRI. NeuroImage. 2007;36(1):144–152.

36. Liu D, Dong Z, Zuo X, Wang J, Zang Y. Eyes-open/eyes-closed dataset sharing for reproducibility evaluation of resting state fMRI data analysis methods. Neuroinformatics. 2013;11(4):469–476.

37. Zou Q. Miao X. Liu D. Wang D. J. Zhuo Y. Gao J. H. Reliability comparison of spontaneous brain activities between BOLD and CBF contrasts in eyes-open and eyes-closed resting states. NeuroImage. 2015;121:91–105.

38. Wang JJ, Chen X, Sah SK, et al. Amplitude of low-frequency fluctuation (ALFF) and fractional ALFF in migraine patients: a resting-state functional MRI study. Clinical Radiology. 2016;71(6):558–564.

39. Zhou F, Huang S, Zhuang Y, Gao L, Gong H. Frequency-dependent changes in local intrinsic oscillations in chronic primary insomnia: a study of the amplitude of low-frequency fluctuations in the resting state. NeuroImage: Clinical. 2017;15:458–465.

40. Yang L, Yan Y, Wang Y, et al. Gradual disturbances of the amplitude of low-frequency fluctuations (ALFF) and fractional ALFF in Alzheimer spectrum. Frontiers in Neuroscience. 2018;12:975.

41. Hoptman MJ, Zuo XN, Butler PD, et al. Amplitude of low-frequency oscillations in schizophrenia: a resting state fMRI study. Schizophrenia Research. 2010;117(1):13–20.

42. Wang L, Dai W, Su Y, et al. Amplitude of low-frequency oscillations in first-episode treatment-naive patients with major depressive disorder: a resting-state functional MRI study. PLOS ONE 2012;7(10):e48658.

43. Zhou G, Liu P, Wang J, et al. Fractional amplitude of low-frequency fluctuation changes in functional dyspepsia: a resting-state fMRI study. Magnetic Resonance Imaging. 2013;31(6):996–1000.

44. Zhan J, Gao L, Zhou F, et al. (2016). Amplitude of low-frequency fluctuations in multiple-frequency bands in acute mild traumatic brain injury. Frontiers in Human Neuroscience. 2016;10:27.

45. Patel P, Al-Dayeh L, Singh M. Localisation of alpha activity by simultaneous fMRI and EEG measurements. Proceedings of the International Society for Magnetic Resonance in Medicine 1997;13(18):2487–2492.

46. Goldman RI, Stern JM, Engel JJ, Cohen MS. Simultaneous EEG and fMRI of the alpha rhythm. NeuroReport. 2002;13(18):2487–2492.

47. Moosmann M, Ritter P, Krastel I, et al. Correlates of alpha rhythm in functional magnetic resonance imaging and near infrared spectroscopy. NeuroImage. 2003;20(1):145–158.

48. Scheeringa R, Petersson KM, Kleinschmidt A, Jensen O, Bastiaansen MC. EEG alpha power modulation of fMRI resting-state connectivity. Brain Connectivity. 2012;2(5):254–264.

49. Klimesch W, Sauseng P, Hanslmayr S. EEG alpha oscillations: the inhibition–timing hypothesis. Brain Research Reviews. 2007;53(1):63–88.

50. Bottari D, Troje NF, Ley P, Hense M, Kekunnaya R, Röder B. Sight restoration after congenital blindness does not reinstate alpha oscillatory activity in humans. Scientific Reports. 2016;6(1):1–10.

51. Novikova LA, eds. Blindness and the electrical activity of the brain: electroencephalographic studies of the effects of sensory impairment. American Foundation for the Blind; 1974.

52. Birbaumer N. Das Elektro-Encephalogramm bei Blindgeborenen. In: Haider M, eds. Neuropsychologie. Bern: Huber; 1971:128–146.

53. Adrian ED, Matthews BH. The Berger rhythm: potential changes from the occipital lobes in man. Brain. 1934;57(4):355–385.

54. Kriegseis A, Hennighausen E, Rösler F, Röder B. Reduced EEG alpha activity over parieto-occipital brain areas in congenitally blind adults. Clinical Neurophysiology. 2006;117(7):1560–1573.

55. Hawellek DJ, Schepers IM, Roeder B, Engel AK, Siegel M, Hipp JF. Altered intrinsic neuronal interactions in the visual cortex of the blind. Journal of Neuroscience. 2013;33(43):17072–17080.

56. Lubinus C, Orpella J, Keitel A, et al. Data-driven classification of spectral profiles reveals brain region-specific plasticity in blindness. Cerebral Cortex. 2020;00:1–18.

57. Wang C, Rajagovindan R, Han SM, Ding M. Top-down control of visual alpha oscillations: sources of control signals and their mechanisms of action. Frontiers in Human Neuroscience. 2016;10:15.

58. Popov T, Kastner S, Jensen O. FEF-controlled alpha delay activity precedes stimulus-induced gamma-band activity in visual cortex. Journal of Neuroscience. 2017;37(15):4117–4127.

59. Schölvinck ML, Maier A, Frank QY, Duyn JH, Leopold DA. Neural basis of global resting-state fMRI activity. Proceedings of the National Academy of Sciences. 2010;107(22):10238–10243.

60. Fiser J, Berkes P, Orban G, Lengyel M. Probabilistic computation: A possible functional role for spontaneous activity in the cortex. Perception. 2011;40:53–53.

61. Guerreiro MJ, Putzar L, Röder B. The effect of early visual deprivation on the neural bases of auditory processing. Journal of Neuroscience. 2016;36(5):1620–1630.

62. Burton H, Snyder AZ, Raichle ME. Resting state functional connectivity in early blind humans. Frontiers in Systems Neuroscience. 2014;8:51.

63. Yan CG, Wang XD, Zuo XN, Zang YF. DPABI: Data Processing & Analysis for (Resting-State) Brain Imaging. Neuroinformatics. 2016;14:339–351.

64. Woletz M, Hoffmann A, Tik M, et al. Beware detrending: Optimal preprocessing pipeline for low-frequency fluctuation analysis. Human Brain Mapping. 2019;40(5):1571–1582.

65. Song XW, Dong ZY, Long XY, et al. REST: a toolkit for resting-state functional magnetic resonance imaging data processing. PLOS ONE. 2011;6(9):e25031.

66. Zhou Z, Wang JB, Zang YF, Pan G. PAIR Comparison between Two Within-Group Conditions of Resting-State fMRI Improves Classification Accuracy. Frontiers in Neuroscience. 2018;11:740.

67. Zuo XN, Di Martino A, Kelly C, et al. The oscillating brain: complex and reliable. NeuroImage, 2010;49(2):1432–1445.

68. Cremers HR, Wager TD, Yarkoni T. The relation between statistical power and inference in fMRI. PLOS ONE. 2017;12(11):e0184923.

69. Greve DN, Fischl B. False positive rates in surface-based anatomical analysis. NeuroImage 2018;171:6–14.

70. Eklund A, Nichols TE, Knutsson H. Cluster failure: Why fMRI inferences for spatial extent have inflated false-positive rates. Proceedings of the National Academy of Sciences. 2016;113(28):7900–7905.

71. Bourgeois JP, Goldman-Rakic PS, Rakic P. Formation elimination and stabilization of synapses in the primate cerebral cortex. In: Ganzzaniga MS, eds. The New Cognitive Neurosciences. MIT Press; 2000;45-54.

72. Anurova I, Renier LA, De Volder AG, Carlson S, Rauschecker JP. Relationship between cortical thickness and functional activation in the early blind. Cerebral Cortex. 2015;25(8):2035–2048.

73. Bridge H, Cowey A, Ragge N, Watkins K. Imaging studies in congenital anophthalmia reveal preservation of brain architecture in ‘visual’ cortex. Brain. 2009;132(12):3467–3480.

74. Jiang J, Zhu W, Shi F. Thick visual cortex in the early blind. Journal of Neuroscience. 2009;29(7):2205–2211.

75. Park HJ, Lee JD, Kim EY, et al. Morphological alterations in the congenital blind based on the analysis of cortical thickness and surface area. NeuroImage. 2009;47(1):98–106.

76. Qin W, Liu Y, Jiang T, Yu C. The development of visual areas depends differently on visual experience. PLOS ONE. 2013;8(1):e53784.

77. Guerreiro, M. J., Erfort, M. V., Henssler, J., Putzar, L., & Röder, B. (2015). Increased visual cortical thickness in sight-recovery individuals. Human Brain Mapping, 36(12), 5265–5274.

78. Hölig C, Guerreiro MJS, Röder B, Shareef I, Kekunnaya R, Lingareddy S. Cortical thickness in sight-recovery and congenitally blind individuals. In: Society for Neuroscience 2019. Chicago; 2019. Program No. 308.03 / L25. 2019 Neuroscience Meeting Planner.

79. Chugani HT. PET scanning studies of human brain development and plasticity. Developmental Neuropsychology. 1999;16(3):379–381.

80. De Volder AG, Bol A, Blin J, et al. Brain energy metabolism in early blind subjects: neural activity in the visual cortex. Brain research. 1997;750(1-2):235–244.

81. Wang XJ. Macroscopic gradients of synaptic excitation and inhibition in the neocortex. Nature Reviews Neuroscience. 2020;21:169–178.

82. Reh RK, Dias BG, Iii CAN. Critical period regulation across multiple timescales. Proceedings of the National Academy of Sciences. 2020;117(38):23242–23251.

83. McGee AW, Yang Y, Fischer QS, Daw NW, Strittmatter SM. Experience-driven plasticity of visual cortex limited by myelin and Nogo receptor. Science. 2005;309(5744):2222–2226.

84. Jensen O, Mazaheri A. Shaping functional architecture by oscillatory alpha activity: Gating by inhibition. Frontiers in Human Neuroscience. 2010;4:e186.

85. Lüchinger R, Michels L, Martin E, Brandeis D. Brain state regulation during normal development: Intrinsic activity fluctuations in simultaneous EEG–fMRI. NeuroImage. 2012;60(2):1426–1439.

86. Ip IB, Emir UE, Parker AJ, Campbell J, Bridge H. Comparison of neurochemical and BOLD signal contrast response functions in the human visual cortex. Journal of Neuroscience. 2019;39(40): 7968–7975.

87. Weaver KE, Richards TL, Saenz M, Petropoulos H, Fine I. Neurochemical changes within human early blind occipital cortex. Neuroscience. 2013;252:222–233.

88. Coullon GS, Emir UE, Fine I, Watkins KE, Bridge H. Neurochemical changes in the pericalcarine cortex in congenital blindness attributable to bilateral anophthalmia. Journal of Neurophysiology. 2015;114(3):1725–1733.

89. Faini G, Aguirre A, Landi S, et al. Perineuronal nets control visual input via thalamic recruitment of cortical PV interneurons. eLife. 2018;7:e41520.

90. Hyvärinen J, Carlson S, Hyvärinen L. Early visual deprivation alters modality of neuronal responses in area 19 of monkey cortex. Neuroscience Letters. 1981;26(3):239–243.

91. Pan P, Zhou Y, Fang F, Zhang G, Ji Y. Visual deprivation modifies oscillatory activity in visual and auditory centers. Animal cells and systems. 2018;22(3):149–156.

92. Kral A, Yusuf PA, Land R. Higher-order auditory areas in congenital deafness: Top-down interactions and corticocortical decoupling. Hearing Research. 2017;343:50–63.

93. Castaldi E, Lunghi C, Morrone MC. Neuroplasticity in adult human visual cortex. Neuroscience & Biobehavioral Reviews. 2020;112:542–552.

94. Hyvärinen J, Hyvärinen L, Linnankoski I. Modification of parietal association cortex and functional blindness after binocular deprivation in young monkeys. Experimental Brain Research. 1981;42(1):1–8.

95. Carlson S, Pertovaara A, Tanila H. Late effects of early binocular visual deprivation on the function of Brodmann’s area 7 of monkeys (Macaca arctoides). Developmental Brain Research. 1987;33(1):101–111.

96. Ley P, Bottari D, Shenoy BH, Kekunnaya R, Röder B. Partial recovery of visual–spatial remapping of touch after restoring vision in a congenitally blind man. Neuropsychologia. 2013;51(6):1119–1123.

97. Azañón E, Camacho K, Morales M, Longo MR. The sensitive period for tactile remapping does not include early infancy. Child Development. 2018;89(4):1394–1404.

98. Xu P, Huang R, Wang J, et al. Different topological organization of human brain functional networks with eyes open versus eyes closed. NeuroImage. 2014;90:246–255.

99. Putzar L, Goerendt I, Heed T, Richard G, Büchel C, Röder B. The neural basis of lip-reading capabilities is altered by early visual deprivation. Neuropsychologia. 2010;48(7):2158–2166.

100. Putzar L, Goerendt I, Lange K, Rösler F, Röder B. Early visual deprivation impairs multisensory interactions in humans. Nature Neuroscience. 2007;10(10):1243–1245.

101. Rohe T. Noppeney U. Reliability-weighted integration of audiovisual signals can be modulated by top-down attention. eNeuro. 2018;5(1).

