## Supplementary Materials for "Typical resting state activity of the brain requires visual input during an early sensitive period"

Biological Psychology and Neuropsychology

University of Hamburg

Von-Melle-Park 11

20146 Hamburg, Germany

**Running title**: Resting state activity in sight recovery

**Supplementary Table 1.** Brain regions showing differences in ALFF in the EO compared to the EC condition in sighted control individuals (SC group. n=28).

| Hemisphere | Brain region | Brodmann area | Cluster size | Peak voxel MNI coordinates | | | Peak voxel t-value |
|---|---|---|---|---|---|---|---|
| | | | | x | y | z | |
| L | Superior Temporal Gyrus / Postcentral Gyrus / Precentral Gyrus | 2, 3, 4, 6 22 | 537 | -52 | -16 | 12 | -4.91 |
| L | Superior Parietal Gyrus / Middle Occipital Gyrus | 7, 19 | 322 | -18 | -68 | 48 | 5.91 |
| R | Middle Occipital Gyrus / Superior Occipital Gyrus / Superior Parietal Gyrus | 7, 19 | 235 | 41 | -78 | 20 | 5.44 |
| R | Postcentral Gyrus / Precentral Gyrus | 2, 3, 4 | 180 | 41 | -6 | 12 | -4.86 |
| L | Precuneus / Cingulate Gyrus | 7, 23 | 151 | -1 | -54 | 24 | 5.11 |
| L | Precentral Gyrus / Superior Frontal Gyrus | 4, 8, 9 | 133 | -18 | -19 | 80 | -4.65 |
| R | Middle Temporal Gyrus / Inferior Temporal Gyrus | 20, 21 | 80 | 48 | -64 | -4 | -5.11 |
| L | Superior Frontal Gyrus / Middle Frontal Gyrus | 8, 9, 46 | 62 | -18 | 56 | 12 | 4.19 |

*Note.* Clusters showing group differences in ALFF between the EO and the EC condition at $p < .01$ voxel-wise and $p < .05$ cluster-wise after Gaussian random field (GRF) correction for multiple comparisons. MNI coordinates and t-values are derived from the peak voxel of the cluster. A positive t-value means greater ALFF in the EO condition and a negative t-value means greater ALFF in the EC condition. EO = eyes open. EC = eyes closed. MNI = Montreal Neurological Institute coordinates system. L = left. R = right.

**Supplementary Table 2.** Brain regions showing differences in ALFF in the EO compared to the EC condition for each group.

| Hemisphere | Brain region | Brodmann areas | Cluster size | Peak voxel MNI coordinates x | y | z | Peak voxel t-value |
|---|---|---|---|---|---|---|---|
| **ALFF in the SC group (n=19) in the EO vs. the EC condition** | | | | | | | |
| L | Postcentral Gyrus / Precentral Gyrus | 2, 3, 4 | 193 | -49 | -23 | 40 | -5.44 |
| L | Medial Frontal Gyrus / Superior Frontal Gyrus / Middle Frontal Gyrus | 8, 9, 46 | 122 | -4 | 53 | 8 | 4.67 |
| R | Supramarginal Gyrus / Postcentral Gyrus / Precentral Gyrus | 2, 3, 4, 40 | 79 | 48 | -26 | 36 | -4.40 |
| R | Middle Temporal Gyrus / Middle Occipital Gyrus / Inferior Temporal Gyrus | 19, 20, 21 | 72 | 48 | -64 | -4 | -5.40 |
| R | Cingulate Gyrus / Medial Frontal Gyrus / Supplementary Motor Area | 6, 8, 9, 23 | 70 | 13 | -13 | 48 | -3.78 |
| R | Medial Frontal Gyrus / Superior Frontal Gyrus | 8, 9 | 57 | 6 | -13 | 80 | -3.71 |
| L | Superior Parietal Gyrus / Precuneus | 7 | 55 | -18 | -68 | 48 | 5.04 |
| L | Precuneus / Cingulate Gyrus | 7, 23 | 49 | -11 | -50 | 24 | 3.40 |
| **ALFF in the CC group (n=19) in the EO vs. the EC condition** | | | | | | | |
| L | Calcarine Gyrus / Lingual Gyrus / Middle Occipital Gyrus | 17, 18, 19 | 359 | -28 | -85 | 16 | 4.62 |
| L | Cingulate Gyrus | 23 | 102 | -4 | -44 | 16 | -4.51 |
| R | Postcentral Gyrus / Supramarginal Gyrus | 2, 3, 40 | 50 | 48 | -30 | 56 | -4.28 |
| **ALFF in the DC group (n=11) in the EO vs. the EC condition** | | | | | | | |
| R | Precentral Gyrus / Postcentral Gyrus / Superior Temporal Gyrus / Supplementary Motor Area / Cingulate Gyrus | 2, 3, 4, 6, 22, 23 | 1018 | 44 | -13 | 16 | -7.20 |

| | | | | | | | |
|---|---|---|---|---|---|---|---|
| R | Middle Frontal Gyrus / Inferior Frontal Gyrus / Superior Frontal Gyrus | 8, 9, 45, 46 | 311 | 44 | 25 | 0 | 6.97 |
| L | Superior Frontal Gyrus | 8, 9 | 58 | -18 | 39 | 48 | 4.38 |
| L | Superior Frontal Gyrus / Middle Frontal Gyrus | 8, 9, 46 | 55 | -18 | 67 | 16 | 5.38 |
| R | Superior Frontal Gyrus / Middle Frontal Gyrus | 8, 9, 46 | 50 | 13 | 39 | 40 | 4.92 |
| R | Middle Temporal Gyrus | 21 | 46 | 41 | -57 | -12 | -5.01 |
| R | Calcarine Gyrus / Lingual Gyrus | 17, 18 | 44 | 23 | -47 | -4 | -5.34 |
| **ALFF in the CB group (n=9) in the EO vs. the EC condition** | | | | | | | |
| L | Superior Frontal Gyrus / Middle Frontal Gyrus / Medial Frontal Gyrus | 8, 9, 46 | 679 | -38 | 60 | 0 | 12.99 |
| L | Precuneus / Superior Parietal Gyrus | 7 | 45 | -18 | -71 | 40 | -5.37 |

Note. Clusters showing group differences in ALFF between the EO and the EC condition at p < .01 voxel-wise and p < .05 cluster-wise after Gaussian random field (GRF) correction for multiple comparisons. MNI coordinates and t-values are derived from the peak voxel of the cluster. A positive t-value means greater ALFF in the EO condition and a negative t-value means greater ALFF in the EC condition. EO = eyes open. EC = eyes closed. MNI = Montreal Neurological Institute coordinates system. L = left. R = right.

**Supplementary Table 3.** Brain regions showing differences in ALFF between the tested groups separately in the EO and in the EC condition.

| Hemisphere | Brain region | Brodmann area | Cluster size | Peak voxel MNI coordinates | | | Peak voxel t-value |
|---|---|---|---|---|---|---|---|
| | | | | x | y | z | |
| **ALFF in the CC group (n=19) vs. SC group (n=19)** | | | | | | | |
| **in the EO condition** | | | | | | | |
| R | Calcarine Gyrus / Lingual Gyrus / Middle Occipital Gyrus / Inferior Occipital Gyrus | 17, 18, 19 | 286 | 30 | -92 | -12 | 4.26 |
| R | Precuneus / Cingulate Gyrus | 7, 23 | 195 | 6 | -54 | 24 | -5.04 |
| R | Inferior Frontal Gyrus / Superior Frontal Gyrus / Middle Frontal Gyrus | 8, 9, 45, 46 | 137 | 27 | 46 | 40 | -4.48 |
| L | Superior Temporal Gyrus / Angular Gyrus / Supramarginal Gyrus | 22, 39, 40 | 126 | -56 | -61 | 24 | -3.78 |
| R | Superior Temporal Gyrus / Angular Gyrus / Supramarginal Gyrus | 22, 39, 40 | 115 | 48 | -54 | 24 | -4.53 |
| **in the EC condition** | | | | | | | |
| R | Middle Frontal Gyrus / Superior Frontal Gyrus | 8, 9, 46 | 153 | 30 | 18 | 48 | -5.05 |
| R | Supramarginal Gyrus / Angular Gyrus / Superior Temporal Gyrus | 22, 39, 40 | 91 | 65 | -50 | 20 | -4.58 |
| **ALFF in the DC group (n=11) vs. SC group (n=11)** | | | | | | | |
| **in the EO condition** | | | | | | | |
| L | Precuneus / Cingulate Gyrus | 7, 23 | 96 | -8 | -61 | 28 | -5.40 |
| **ALFF in the CB group (n=9) vs. SC group (n=9)** | | | | | | | |
| **in the EO condition** | | | | | | | |
| L | Superior Frontal Gyrus / Medial Frontal Gyrus | 8, 9 | 104 | -32 | 63 | 0 | 5.03 |
| L | Middle Occipital Gyrus / Fusiform Gyrus | 19, 37 | 51 | -45 | -54 | -20 | 3.96 |
| **in the EC condition** | | | | | | | |

| | | | | | | | |
|---|---|---|---|---|---|---|---|
| L | Middle Occipital Gyrus | 19 | 103 | -35 | -78 | 16 | 4.97 |
| R | Precuneus / Superior Occipital Gyrus / Middle Occipital Gyrus | 7, 19 | 79 | 30 | -71 | 36 | 6.01 |
| R | Lingual Gyrus / Fusiform Gyrus | 18, 37 | 68 | 23 | -64 | -8 | 4.78 |
| R | Inferior Frontal Gyrus / Superior Temporal Gyrus | 22, 45 | 65 | 54 | 18 | 0 | -5.49 |
| L | Fusiform Gyrus | 37 | 62 | -35 | -54 | -12 | 5.51 |
| R | Middle Frontal Gyrus / Inferior Frontal Gyrus | 45, 46 | 58 | 44 | 49 | 12 | -4.00 |

Note. Clusters showing group differences in ALFF between the tested groups separately in the EO and in the EC condition at $p < .01$ voxel-wise and $p < .05$ cluster-wise after Gaussian random field (GRF) correction for multiple comparisons. MNI coordinates and t-values are derived from the peak voxel of the cluster. A positive t-value means greater ALFF in the CC group, the DC group and the CB group, respectively, and a negative t-value means greater ALFF in the SC group. EO = eyes open. EC = eyes closed. MNI = Montreal Neurological Institute coordinates system. L = left. R = right.

**Supplementary Table 4.** Brain regions showing differences in ALFF between the CC individuals (n=19) and CB individuals (n=9) separately in the EO and in the EC condition. .

| Hemisphere | Brain region | Brodmann area | Cluster size | Peak voxel MNI coordinates | | | Peak voxel t-value |
|---|---|---|---|---|---|---|---|
| | | | | x | y | z | |
| **in the EO condition** | | | | | | | |
| L | Superior Frontal Gyrus [a] | 8, 9 | 154 | -8 | 1 | 72 | 3.48 |
| L | Supramarginal Gyrus / Angular Gyrus | 39, 40 | 76 | -56 | -54 | 28 | -4.35 |
| R | Supramarginal Gyrus | 40 | 65 | 61 | -40 | 32 | -3.75 |
| **in the EC condition** | | | | | | | |
| R | Calcarine Gyrus / Lingual Gyrus / Middle Occipital Gyrus / Fusiform Gyrus | 17, 18, 19, 37 | 221 | 30 | -71 | -4 | -4.95 |
| L | Middle Occipital Gyrus / Fusiform Gyrus / Cerebellum | 19, 37 | 157 | -35 | -64 | -20 | -6.21 |
| L | Calcarine Gyrus / Lingual Gyrus / Middle Occipital Gyrus | 17, 18, 19 | 132 | -35 | -81 | 16 | -4.48 |

*Note.* Clusters showing group differences in ALFF between the CC individuals (n=19) and CB individuals (n=9) in the EC condition at $p < .01$ voxel-wise and $p < .05$ cluster-wise after Gaussian random field (GRF) correction for multiple comparisons. MNI coordinates and t-values are derived from the peak voxel of the cluster. A positive t-value means greater ALFF in the CC group and a negative t-value means greater ALFF in the CB group. EO = eyes open. EC = eyes closed. MNI = Montreal Neurological Institute coordinates system. L = left. R = right.
[a] This cluster is not visible in the Supplementary Figure 4A.

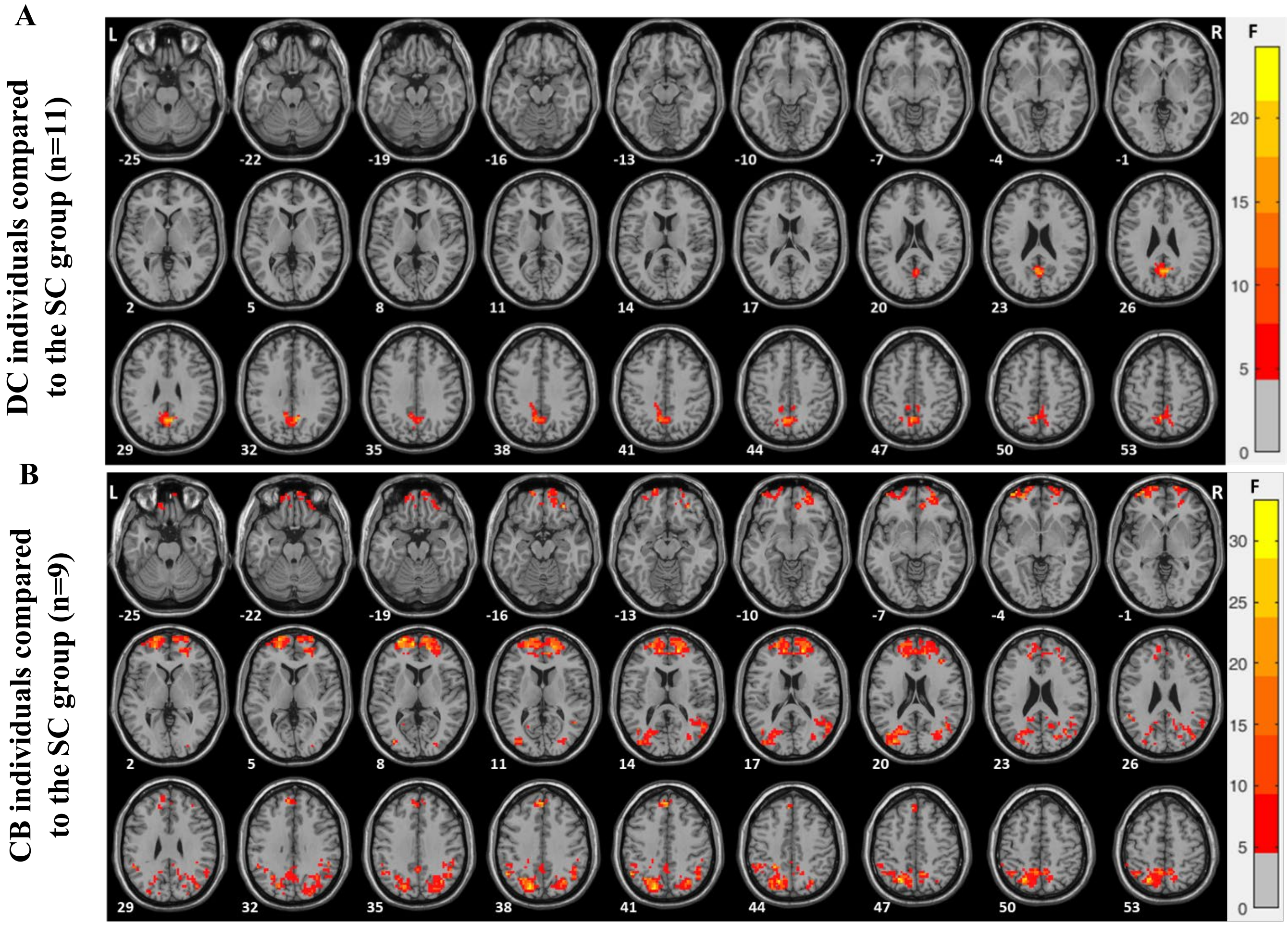


**Supplementary Figure 1 Group differences in ALFF in the EO compared to the EC condition.** Using standardized ALFF (amplitude of low frequency fluctuations) i.e. Z-scores, a mixed 2x2 model (group x condition) was carried out for **(A)** developmental cataract-reversal individuals (DC) vs. SC group (n=11) and **(B)** congenitally blind individuals (CB) vs. SC group (n=9). Regions with significant interaction effects are shown after Gaussian random field theory (GRF) correction for multiple spatial comparisons (voxel-wise $p < .05$, cluster-wise $p < .025$, corrected).

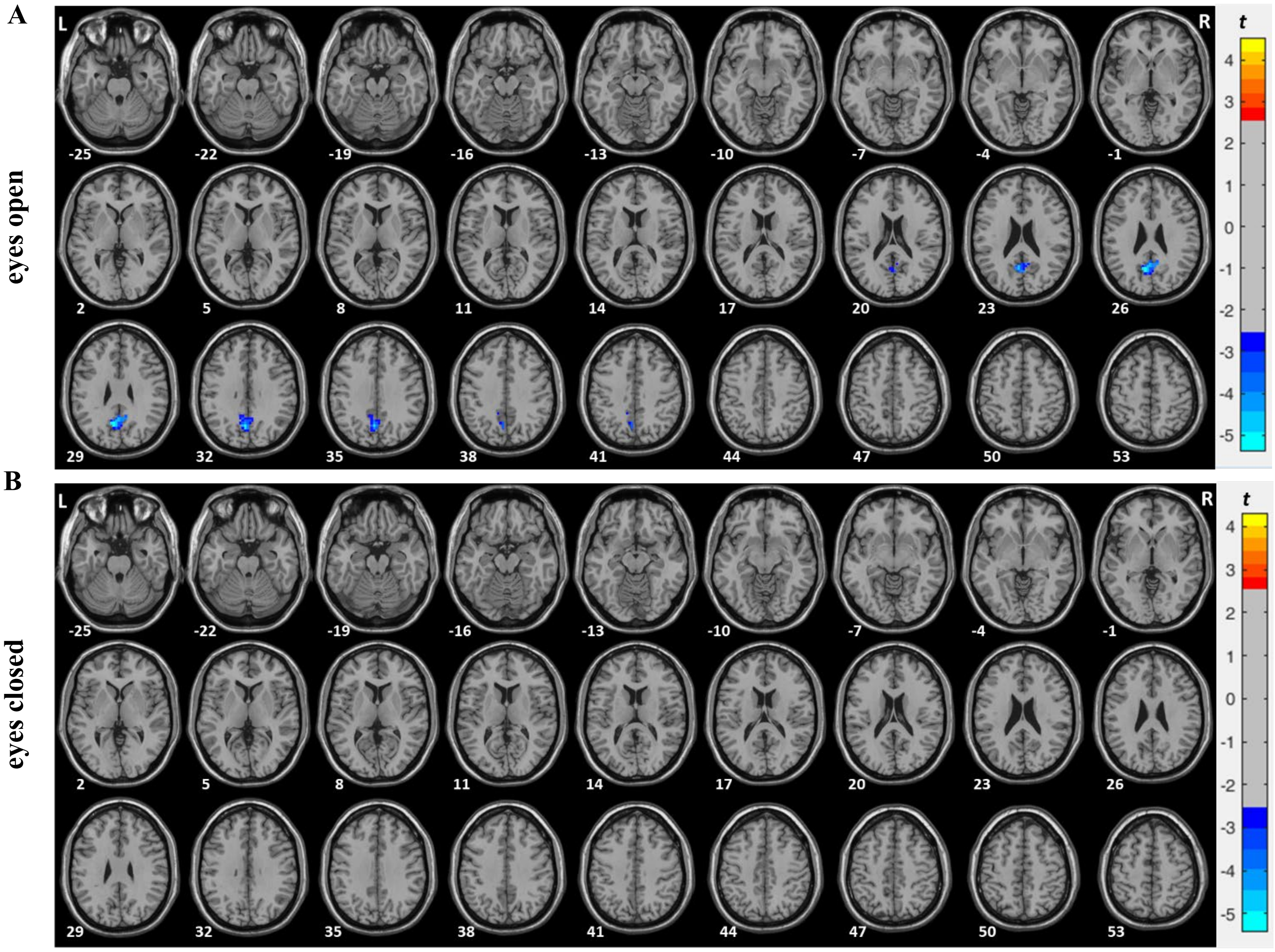


**Supplementary Figure 2 ALFF in the DC group (n=11) compared to the SC group (n=11) in the EO and in the EC condition**. Two-sample t-test results of the amplitude of low frequency fluctuations (ALFF) comparing developmental cataract-reversal individuals (DC) and the SC group (n=11) matched in age and sex to the DC group separately in the **(A)** eyes open (EO) condition and in the **(B)** eyes closed (EC) condition. The red colors denote voxels with significantly higher amplitude for the DC compared to the SC group and the blue colors denote voxels with significantly lower amplitude for the DC compared to the SC group. Significant clusters are shown after Gaussian random field theory (GRF) correction for multiple spatial comparisons (voxel-wise $p < .01$, cluster-wise $p < .05$, corrected).

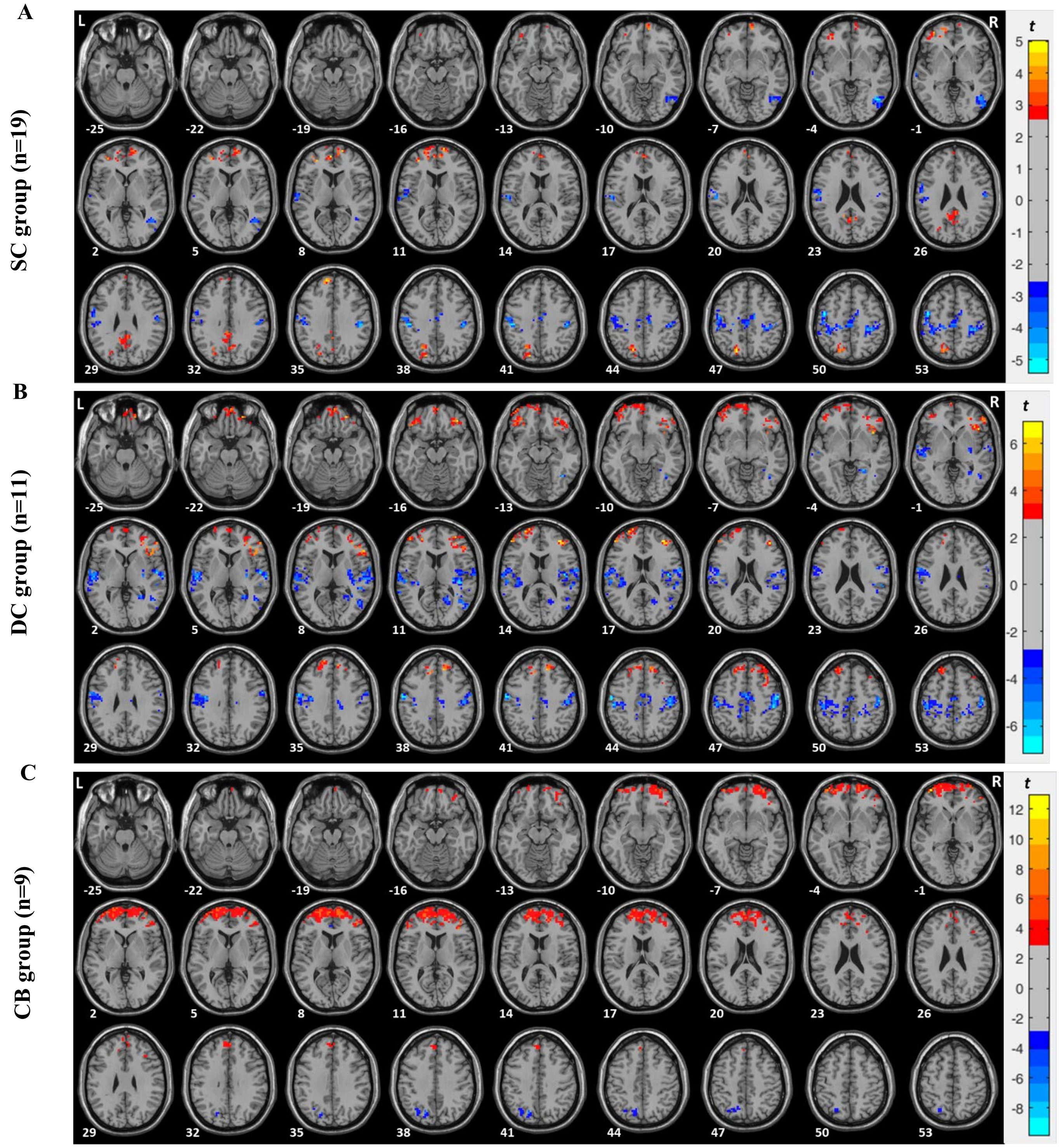


**Supplementary Figure 3 ALFF in the EO compared to the EC condition** Paired t-test results of the amplitude of low frequency fluctuations (ALFF) comparing the eyes open (EO) and the eyes closed (EC) condition in the group of **(A)** sighted individuals (n=19) matched in age and sex to CC individuals **(B)** developmental cataract-reversal individuals and **(C)** congenitally blind individuals. The red colors denote voxels with significantly higher amplitude for the EO compared to the EC condition and the blue colors denote voxels with significantly lower amplitude for the EO compared to the EC condition. Significant clusters are shown after Gaussian random field (GRF) correction for multiple spatial comparisons (voxel-wise $p < .01$, cluster-wise $p < .05$, corrected).

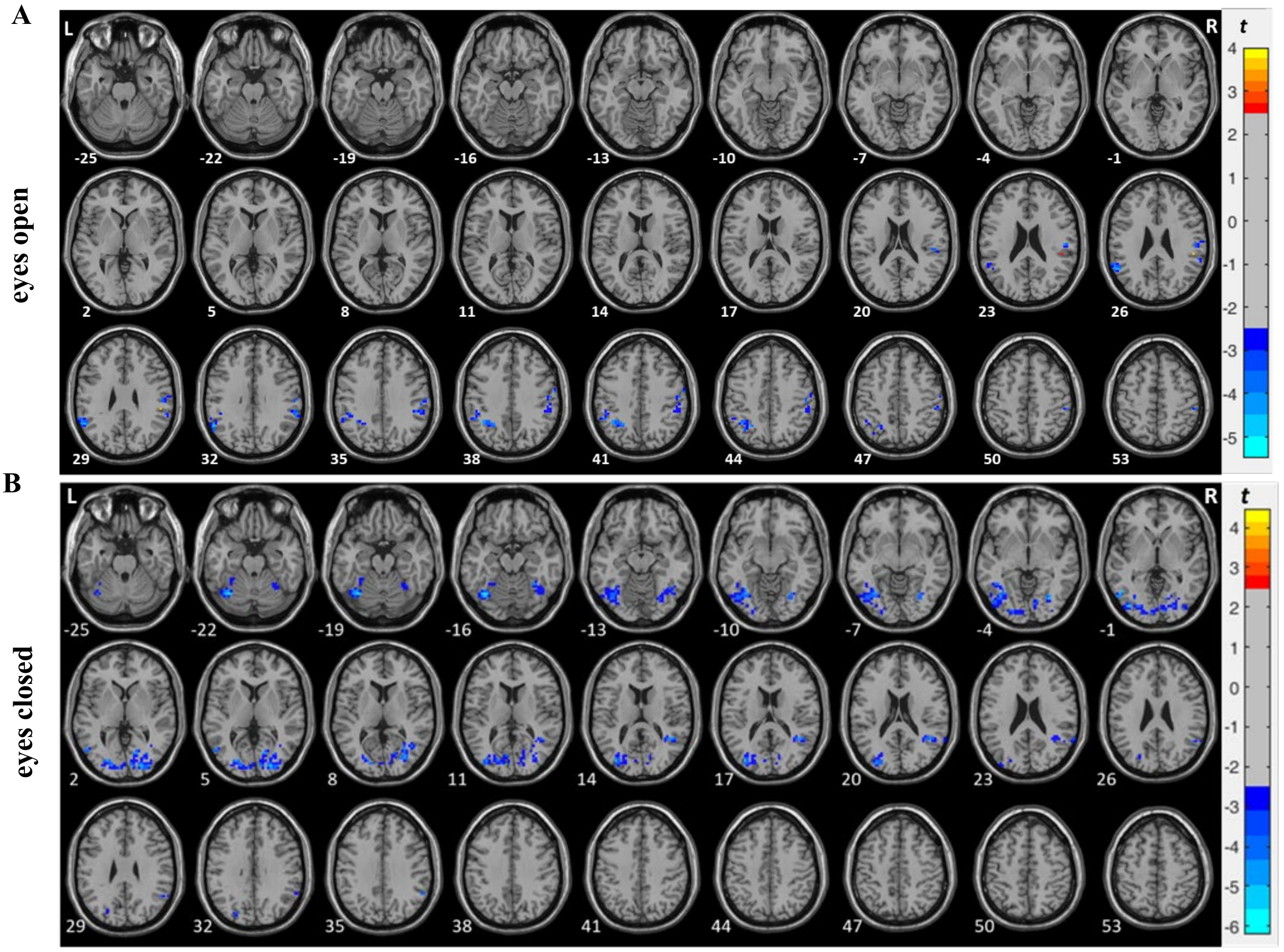


**Supplementary Figure 4 ALFF in the CC group (n=19) compared to the CB group (n=9) in the EO and in the EC condition**. Two-sample t-test results of the amplitude of low frequency fluctuations (ALFF) comparing congenital cataract-reversal individuals (CC group, n=19) and congenitally blind individuals (CB group, n=9) in the **(A)** eyes open (EO) and in the **(B)** eyes closed (EC) condition. The red colors denote voxels with significantly higher amplitude for the CC group compared to the CB group and the blue colors denote voxels with significantly lower amplitude for the CC group compared to the CB group. Significant clusters are shown after Gaussian random field theory (GRF) correction for multiple spatial comparisons (voxel-wise $p < .01$, cluster-wise $p < .05$, corrected).
